# How do biomarkers dance? Specific moves of defense and damage biomarkers for biological interpretation of dose-response model trends

**DOI:** 10.1101/2023.08.04.551999

**Authors:** Simon Colas, Séverine Le Faucheur

## Abstract

Omics and multi-omics studies are currently increasingly used in ecotoxicology to highlight the induction of known or new biomarkers when an organism is exposed to one (or more) contaminant(s). Although it is virtually impossible to identify all biomarkers from all possible organisms, biomarkers can be grouped into two categories, defense or damage biomarkers and they have a limited number of response trends. Our working hypothesis is that defense and damage biomarkers show different dose-response patterns. A meta-analysis of 156 articles and 2,595 observations of dose-response curves of well-known defense and damage biomarkers was carried out in order to characterize the response trends of these biological parameters in a large panel of living organisms (18 phyla) exposed to a wide variety of inorganic or organic contaminants. Defense biomarkers describe biphasic responses (bell-shaped and U-shaped) to a greater extent than damage biomarkers. In contrast, damage biomarkers varied mainly monotonically (decreasing or increasing). Neither the nature of the contaminant nor the type of organisms, whatever the kingdom (Plantae, Animalia, Chromista or Bacteria), influence these specific responses. This result suggests that cellular defense and damage mechanisms are not specific to stressors and are conserved throughout life. The meta-analysis results confirm the usefulness of trend analysis in dose-response models as a biological interpretation of biomarkers in large dataset and their application in determining the concentration ranges inducing defense responses (CRIDeR) and the concentration ranges inducing damage responses (CRIDaR) regardless of the contaminant tested or the organism studied.

**Highlights:**

- We interpreted 2,595 biomarker dose-response curves generated by chemical exposure.
- Defense biomarkers mainly describe biphasic (bell- or U-shaped) trends.
- Damage biomarkers mainly describe monotonic (decreasing or increasing) trends.
- Cellular defense and damage responses appear to have been conserved during evolution.
- Response trend analysis is a promising tool for environmental risk assessment.

**Graphical abstract:** 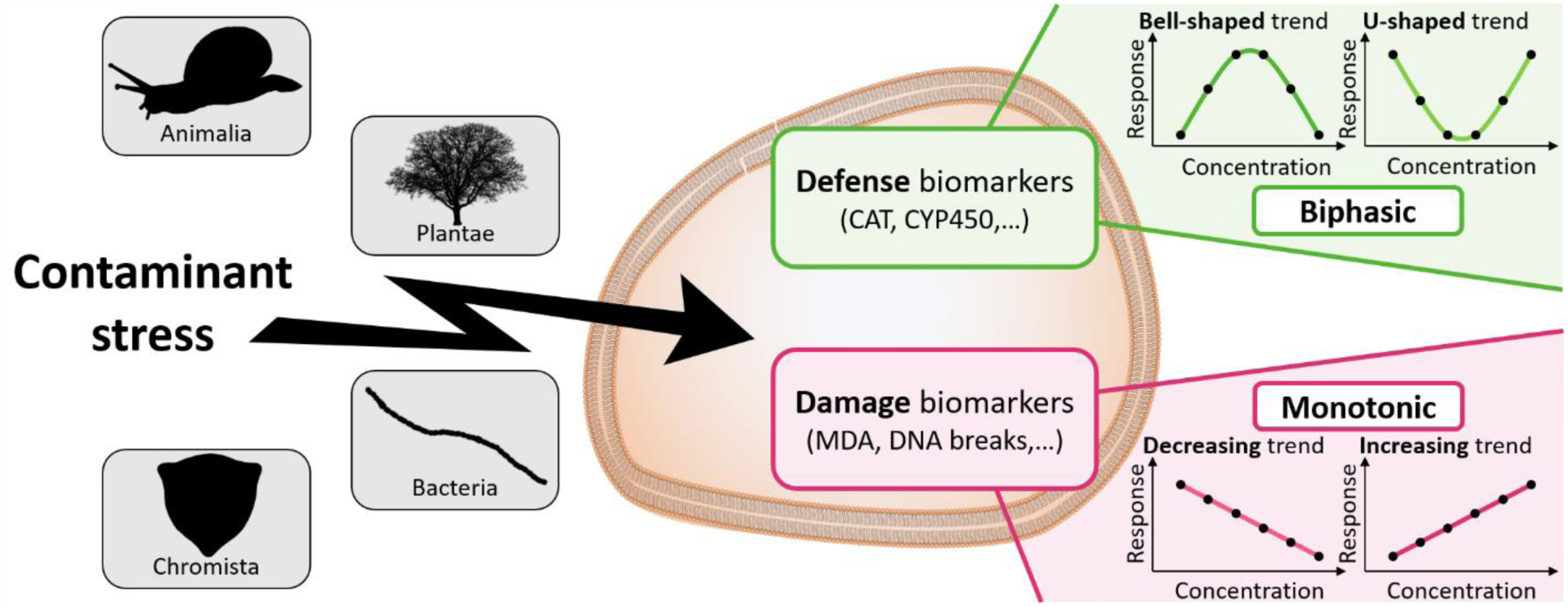

## 1. Introduction

Lethal endpoints are most commonly used to build dose-response curves to feed in risk assessments of contaminants. There is however a growing interest in identifying sublethal endpoints that can provide information on early effects at the individual level. Biomarkers can be used as indicators of sublethal effects, to quantify a stress on an organism of interest. They can be also used for environmental monitoring. A biomarker is defined as an observable and/or measurable change at the molecular, biochemical, cellular, physiological or behavioral level, which indicates the present or past exposure of an organism to at least one stressor[1,2]. It can also provide information on early effects at the individual level[3]. The biological unity of the living world means that the same types of biomarkers can be used for organisms belonging to different taxonomic groups. Well-known biomarkers include catalase (CAT)[4], superoxide dismutase (SOD)[5], heat-shock proteins (HSP)[6] and single-strand DNA breaks[7]. A further distinction can be made between defense biomarkers and damage biomarkers[8]. Defense biomarkers result from protective reactions of the organism exposed to a contaminant. Induction of these mechanisms will allow the organism to cope with a stress. Damage biomarkers, on the other hand, reflect a direct and harmful change caused to the exposed organism. They are the result of an alteration that can lead to an inability to grow, reproduce or even survive.

The identification of new biomarkers is an important field of research today in a context of the increasing complexity of stress caused by global changes[9]. The development of “omics” tools over the last decade is a promising avenue to address that challenge[10–12]. However, despite the recent progress, it is still difficult to provide a biological interpretation to an observation of change in specific sub-cellular or cellular parameters. This is particularly true for metabolomics, for which, a large proportion of the metabolites observed cannot be identified with the current information available in databases. In mass spectrometry, especially with liquid chromatography-mass spectrometry (LC-MS) analyses, only 4%-5% of the peaks can be identified in biological samples[13]. This gap in the reference databases is also encountered in genomics[14] and proteomics[15].

However, even if a biomarker cannot be characterized, it is still possible to model its response as a function of the dose of exposure. These dose-response curves are usually drawn following exposure to increasing concentrations of a substance in order to characterize its toxicity[16]. Four main types of trends can then be observed[17]. In bell-shaped curves, an increasing biomarker response up to a maximum is observed at low doses, followed by a decrease at higher concentrations of the exposure substance. The U-shaped curves follow the same principle but in the opposite direction, *i.e.,* a decrease down to a minimum followed by an increase. These two types of trends can be grouped together as biphasic responses. In contrast, monotonic responses are characterized by continuously increasing or decreasing trends. The increasing curves correspond to a continuous increase of the biological response studied as a function of the increase of the concentrations of the substance of interest. Conversely, the decreasing curves indicate a decrease in the biomarker response as a function of the increase in exposure concentrations.

A very recent study focused on metabolomic response of biofilms exposed to cobalt[18]. The metabolite dose-response curves highlighted a concentration range inducing defense responses (CRIDeR), mainly characterized by bell and U-shape trends, and a concentration range inducing damage responses (CRIDaR) mainly composed of increasing and decreasing trends. These two distinctive ranges of concentrations were validated with the use of biomass and chlorophyll measurements, confirming the usefulness of this novel approach. Our working hypothesis is then that the trends in dose-response curves depend on the role of the biomarker, *i.e.*, defense or damage biomarkers, which can be distinguished according to whether they describe a biphasic or monotonic response. To this end, we conducted a meta-analysis in which we examined the trends of biomarker dose-response curves in several phyla exposed to a contaminant (organic or inorganic). We tested whether this hypothesis was valid regardless of the studied contaminant and phylum. There was no *a priori* on the species in order to have a broad representation of the response to contamination. The conclusions of this study could be used to interpret the dose-response curves plotted from a large data set of unidentified “omics” data and support their use in environmental risk assessment.

## 2. Material and methods

### 2.1. Biomarker compilation

Compilation of biomarkers was carried out searching *Google Scholar* and *Scopus* using combination of keywords, *e.g., biomarker* AND *ecotoxicology* AND *dose-response*. Biomarkers were classified either as defense or damage according to their role in organisms (Table 1 and Table S1)[19,20]. The search was performed between September 2022 and June 2023, without any restriction on the publication date. The aim was to reach an equal proportion of observations of biomarkers of damage and defense (around 1,000 observations for each category), as well as between organic and inorganic contaminants. To achieve this, we had to refine the article search by adding keywords to the initial search: (i) *damage biomarker* OR *defense biomarker* AND *ecotoxicology* AND *dose-response*, (ii) *biomarker* AND *ecotoxicology* AND *dose-response* AND *metal*.

**Table 1:** Examples of biomarker classification (defense and damage) as a function of their molecular, sub-cellular and cellular role.

| Class of biomarkers | Roles | Biomarkers | Reference |
| --- | --- | --- | --- |
| Defense biomarkers | Biotransformation enzymes and associated compounds | Cytochrome P450 (CYP450) | [33]Stegeman et al. (1992) |
|  |  | Glutathione S-transferase (GST) | [34]George (1994) |
|  |  | Glutathione (GSH) | [34]George (1994) |
|  | Toxic efflux | P-glycoproteins | [35]Bard (2000) |
|  |  | Multixenobiotic resistance systems (MXR) | [35]Bard (2000) |
|  | Metals detoxication | Metallothioneins | [36]Hall (2002) |
|  |  | Phytochelatins | [36]Hall (2002) |
|  | Oxidative stress response enzymes and molecules | Superoxide dismutase (SOD) | [33]Stegeman et al. (1992) |
|  |  | Catalase (CAT) | [33]Stegeman et al. (1992) |
|  |  | Glutathione peroxidase (GPx) | [37]Lauterburg et al. (1982) |
|  |  | Glutathione reductase (GR) | [19]Van der Oost et al. (2003) |
|  | Chaperone proteins | Heat shock proteins (HSP) | [38]Feder and Hofmann (1999) |
| Damage biomarkers | Molecular damages | Acetylcholinesterase (AChE) | [39]Payne et al. (1996) |
|  |  | Vitellogenin | [40]Matozzo et al. (2008) |
|  |  | Malondialdehyde (MDA) | [41]Janero (1990) |
|  |  | Photosynthetic pigments (chlorophyll *a/b*, carotenoids) | [42]Pikula et al. (2019) |
|  | DNA damages | DNA adducts | [43]La and Swenberg (1996) |
|  |  | Single-strand DNA breaks | [7]Dhawan and Bajpayee (2009) |
|  |  | Double-strand DNA breaks | [44]Heddle et al. (1983) |
|  | Subcellular and cellular damages | Lysosomal membrane stability | [45]Lowe and Pipe (1994) |
|  | Energetic metabolism damages | Adenylate energy charge (CEA) | [46]Lagadic et al. (1994) |
|  | ROS production | Hydrogen peroxide | [47]Halliwell and Gutteridge (1989) |
|  |  | Superoxide radical | [47]Halliwell and Gutteridge (1989) |
|  |  | Hydroxyl radical | [47]Halliwell and Gutteridge (1989) |

### 2.2. Selection criteria

Once the article compilation was completed, selection criteria were applied after a more detailed analysis of the articles: i) only articles on whole organisms (no cell lines) were considered, without selection criteria on the phylum and the species studied, ii) only experiments performed under controlled conditions were kept (no field studies) to have results represented as dose-response curves, iii) only studies on the toxicity of organic and inorganic contaminants were chosen and iv) the number of tested contaminant concentrations had to be equal to or greater than four to be able to identify a trend in the dose-response studied.

### 2.3. Data extraction

For all articles meeting the selection criteria, the information used for the meta-analysis was extracted as follows. First, each biomarker was classified in either as defense or damage categories according to its biological role (Table 1). The studied species and its taxa were documented (Table S1). Information on the contaminant was also indicated *e.g.* its name, its category (organic or inorganic), the range of concentrations tested, the number of concentrations tested and the exposure time. Finally, the trends of fluctuations in the response of the biomarkers were examined according to the significance of the statistical tests used by the different authors. Two scenarios were encountered.

In the first one, statistical tests were performed to compare the biological response at each tested concentration. If at the highest concentration, the response of the biomarker was significantly higher than that of the control and that no intermediate concentration had a response significantly higher than the response at the highest concentration, then the response trend was characterized as “increasing”. Conversely, if at the highest concentration tested, the response of the biomarker was significantly lower than that of the control and that no intermediate concentration had a response significantly lower than the response at the highest concentrations, then the response trend was characterized as “decreasing”. On the other hand, if at an intermediate concentration (or several if they followed one another), the response of the biomarker studied was both significantly higher than that of the control group and that of the highest concentration tested, then the trend was considered to be “bell-shaped”. In the same way, if at an intermediate concentration (or several if they followed one another), the response of the biomarker studied was both significantly lower than that of the control group and that of the highest concentration, then the trend was considered to be “U-shaped”.

In the second case, the statistical tests compared the biological response only to the control group. Then the trends were determined as follows: the trend was characterized as “increasing” when (i) only the biomarker response at the highest concentration was significantly higher than that of the control or (ii) the biomarker response at the highest concentration was significantly higher than that of the control. All the lower limits of the standard deviations of the response at the previous intermediate concentrations, which were significantly higher than that of the control, had to be lower than the upper standard deviation of the highest concentration. In addition, no response at any intermediate concentrations was to be significantly lower than that of the control. Conversely, the trend was characterized as “decreasing” when (i) only the biomarker response at the highest concentration was significantly lower than that of the control or (ii) the biomarker response at the highest concentration was significantly lower than that of the control. All the upper limits of the standard deviations at the previous intermediate concentrations, which were significantly lower than that of the control, had to be greater than the lower standard deviation of the highest concentration. In addition, no response at any intermediate concentrations was to be significantly higher than that of the control. The trend was characterized as “bell-shaped” when (i) the response of the biomarker at an intermediate concentration (or several if they followed one another) was significantly greater than that of the control group, and (ii) the lower limit of the standard deviation of the response at the intermediate concentration was greater than the upper limit of the standard deviation of the response of the group exposed to the highest concentration. In the same way, when (i) the response of the biomarker at an intermediate concentration (or several if they followed each other) was significantly smaller than that of the control group, and (ii) its upper limit of the standard deviation was smaller than the lower limit of the standard deviation of the response at the highest concentration, then the trend was characterized as “U-shaped”.

In cases where there were no significant response differences between the tested concentrations, then the trend was characterized as “constant”. In all other cases, for example when the response seemed to show more than two trends, it could not be characterized and was tagged as “non-determined” (N.D.). These last two categories of trends (constant and N.D.) were not considered in the meta-analysis.

### 2.4. Data analysis

First, the effect of biomarker type on response trends was investigated. To that end, multinomial logit models were fitted to model the proportion of response change as groups of monotonic or biphasic trends in relation to the biomarker type -damage as reference or defense- (“Effect”), the contaminant studied -inorganic as reference or organic- (“Contaminant”) and the number of studied concentrations (“scale(Concentration)”). Random effect variables were introduced into the basic model, including the bibliographic references, *e.g.*, selected articles, (“Ref”), exposure time (“Time_exposure”) and phylum (“Phylum”). Each combination of random effect variables was tested and the model with the best AIC (Akaike Information Criterion) was chosen[21]. Secondly, a multinomial logit model was also fitted to model the proportion of response trends (divided into four categories: bell-shaped, decreasing, increasing and U-shaped) in relation to the biomarker type (damage as reference or defense), the contaminant studied (inorganic as reference or organic). As before, random effect variables were added to the basic model, including the bibliographic reference, exposure time and phylum information. Each combination of random effect variables was also tested and the model with the best AIC was chosen.

Finally, a multinomial logit model was also fitted to study the type of response (monotonic or biphasic trends) of the most represented phyla (at least 50 observations) along four kingdoms (Animalia, Plantae, Chromista and Bacteria) in the analysis, *i.e.*, Annelida (552), Arthropoda (191), Chlorophyta (323), Chordata (283), Cyanobacteria (51), Magnoliophyta (688), Mollusca (278) and Ochrophyta (97), in relation to the biomarker type (damage as reference or defense), the contaminant studied (inorganic as reference or organic) and the number of studied concentrations with “bibliographic reference” as a random effect variable. Multinomial logit models were fitted using the function *mblogit* in the *mclogit* package (0.9.7.)[22] with the maximum likelihood (ML) as an estimator of variance components and the Penalized Quasi-Likelihood (PQL) method for modeling the random effects set. Statistical analyses were performed on R software.

### 2.5. Bias and *a posteriori* check

The authors developed the strategy for article selection and subsequent data extraction. One author carried out the selection and data extraction. In order to assess the possible interpretation bias in the data extraction, *i.e.*, in the determination of the trend in biological responses, 20% of randomly selected articles (34/156) was additionally analyzed by the second author. A percentage of similarity was calculated between the author 1’s and the author 2’s data extractions, and corrections were made where necessary.

## 3. Results

### 3.1. Data compilation

The process for selecting studies and selected extracted data observations (identification, screening, eligibility and inclusion in the meta-analysis) is reported in a flow diagram (Fig. S1). The global build-up database of the meta-analysis is available in the supplementary information section (Table S1). On 4,126 identified articles, a total of 156 articles were selected for this meta-analysis and 3,176 observations of variation of biomarkers could be extracted. Among them, 581 categorized as “constant” or “N.D.” were subsequently removed. One hundred and nineteen species are represented along 18 phyla (Annelida: 552 observations, Arthropoda: 191, Ascomycota: 6, Cercozoa: 4, Charophyta: 32, Chlorophyta: 332, Chordata: 283, Ciliophora: 16, Cnidaria: 7, Cyanobacteria: 51, Echinodermata: 4, Euglenozoa: 3, Haptophyta: 24, Mollusca: 278, Myzozoa: 6, Ochrophyta: 97, Porifera: 6, Magnoliophyta: 688) and communities (biofilm: 15). The impacts of 176 contaminants (or contaminant combinations) including personal care products, pesticides, herbicides, nanoparticles or metals were studied (Table S1).

### 3.2. Data interpretation

In total, 20% of the total observations were re-extracted (636/3,176) out of the 34 randomly selected articles for comparison of trend analysis. The percentage of data extraction similarity between the two investigators was 97.6%.

### 3.3. Model selection

Based on the AIC values (equations 5 and 13 in Table S2), the best-fit models to explain the variations in the biphasic (bell-shaped and U-shaped) and monotonic (decreasing and increasing) trends were for the fixed effects: the type of biomarker (“Effect”), the type of contaminant (“Contaminant”) and the number of concentrations tested (“scale(Concentration)”) and, for the random effects, the article selected (“Ref”).

### 3.4. Response trends of biomarkers according to their category: defense or damage

In the compiled database, biomarkers distribute almost equally between defense biomarkers (1,325) and damage biomarkers (1,270) (Table S1). Among the defense biomarkers, 402 (30.3%) had a bell-shaped trend, 299 (22.6%) had a decreasing trend, 529 (39.9%) had an increasing trend and 95 (7.2%) had a U-shaped trend. This distribution was different for damage biomarkers, i.e., 139 (10.9%) had a bell-shaped trend, 439 (34.6%) had a decreasing trend, 644 (50.7%) had an increasing trend and 48 (3.8%) had a U-shaped trend (Fig. 1A and Table 2). The proportion of biomarkers with a biphasic trend (bell-shaped or U-shaped) is significantly higher for defense biomarkers than for damage biomarkers (Table 2) (*p-value* < 2e-16) whereas the proportion of biomarkers with a monotonic trend (decreasing or increasing) is significantly greater for damage biomarkers than for defense biomarkers (Table 2) (*p-value* < 2e-16). More precisely, the proportion of bell-shaped trends is significantly greater than that of decreasing and increasing trends for defense biomarkers (Tables 3A-C). For clarity, here and in the rest of the text, *p-values* of the models are only given in their corresponding tables. Conversely, the proportions of decreasing and increasing trends are significantly greater than the bell-shaped trends for damage biomarkers (Tables 3A-C). The same observation was made for the proportions of U-shaped trends compared with those of decreasing and increasing trends (Tables 3B-D).

**Figure 1:**
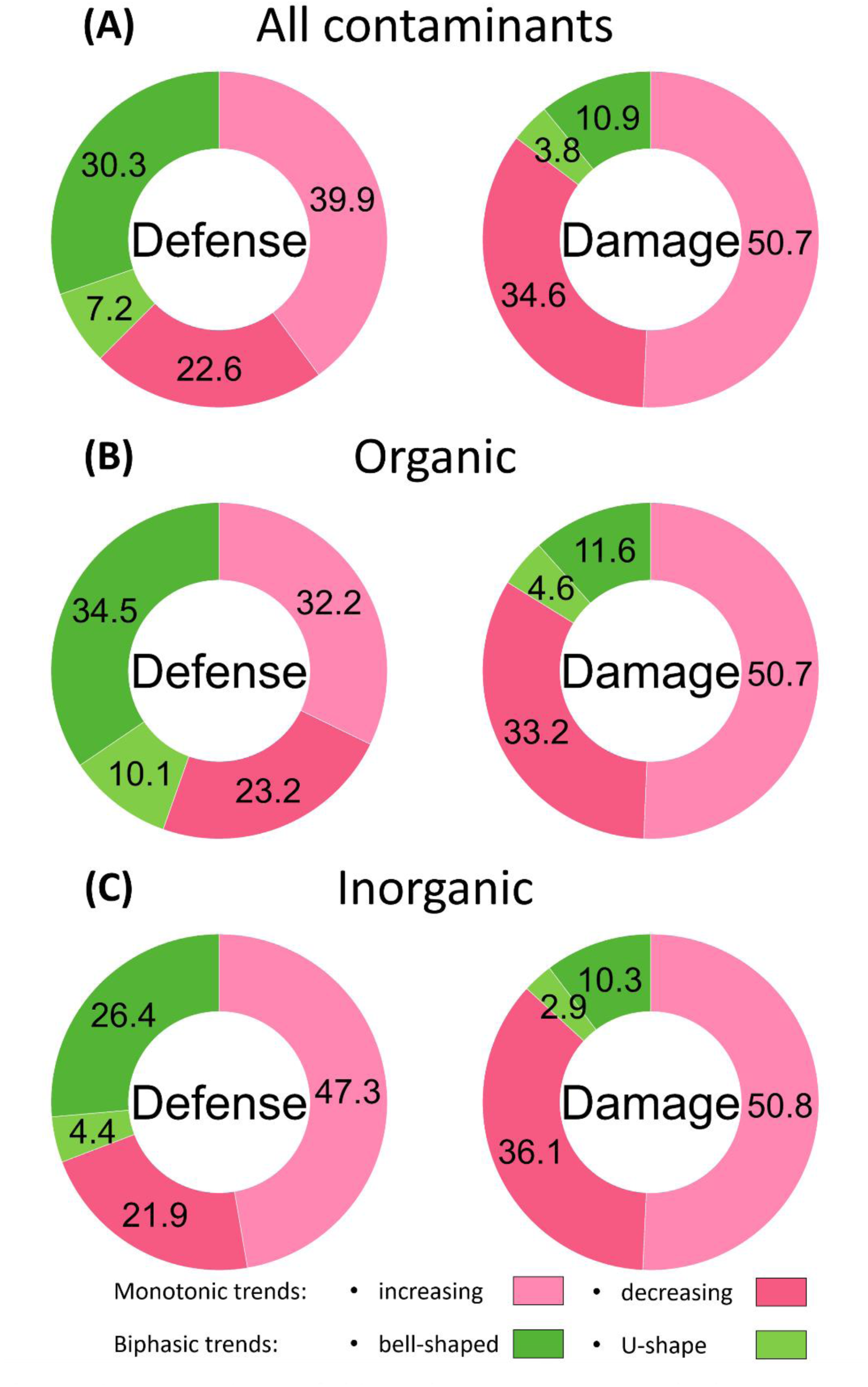
Proportions of biomarker response trends by response mechanism (defense or damage). (A) Influence of the category of biomarkers on the response trends to a contaminant exposure; (B) to an organic contaminant exposure; (C) to an inorganic contaminant exposure.

**Table 2:** Results of the Multinomial Logistic Regression to examine the effect of the type of biomarkers (Defense vs. Damage), the type of contaminants (Inorganic vs. Organic) and the number of tested concentrations on response trends (Biphasic or monotonic). Significant fixed effects are depicted in bold font and indicated with asterisks as follows: * *p* < 0.5, ** *p* < 0.01, *** *p* < 0.001.

| Fixed Effects |  |  |  |  |  |
| --- | --- | --- | --- | --- | --- |
| Contrast | Effect | Estimate | Std.Error | z | p |
| monotonic vs biphasic | Intercept | 2.1344 | 0.1818 | 11.739 | **< 2e-16 \*\*\*** |
|  | Effect-Damage (ref) |  |  |  |  |
|  | Effect-Defense | -1.3108 | 0.1194 | -10.980 | **<2e-16 \*\*\*** |
|  | Contaminant-Inorganic (ref) |  |  |  |  |
|  | Contaminant-Organic | -0.3764 | 0.2287 | -1.646 | 0.0997 |
|  | scale(Concentration) | -0.4664 | 0.1067 | -4.370 | **1.24e-05 \*\*\*** |
| Random Effects |  |  |  |  |  |
| Ref | ~1 |  | Approximate residual deviance: | 2,471 |  |
|  | Estimate | Std. Error | Number of Fisher scoring iterations: | 6 |  |
| biphasic~1 | 0.7488 | 0.05249 | Number of observations | Groups by references: | 156 |
| monotonic ~1 | 0.613 | 0.02514 |  | Individual observations: | 2,595 |
| Model performance |  |  |  |  |  |
| AIC | BIC | RMSE | Sigma |  |  |
| 2,483.06 | 2,518.229 | 0.37 | 0.977 |  |  |

**Table 3:**
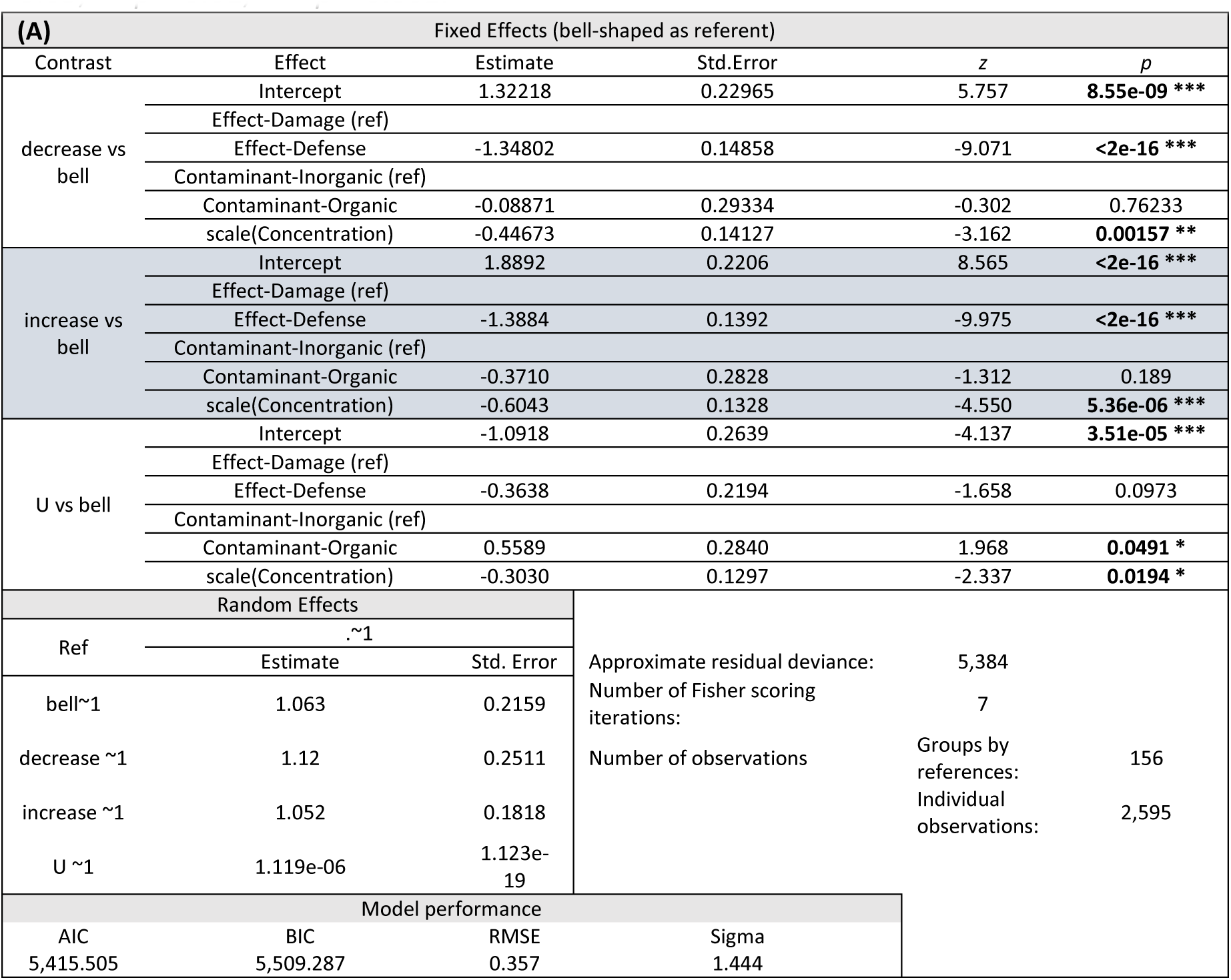
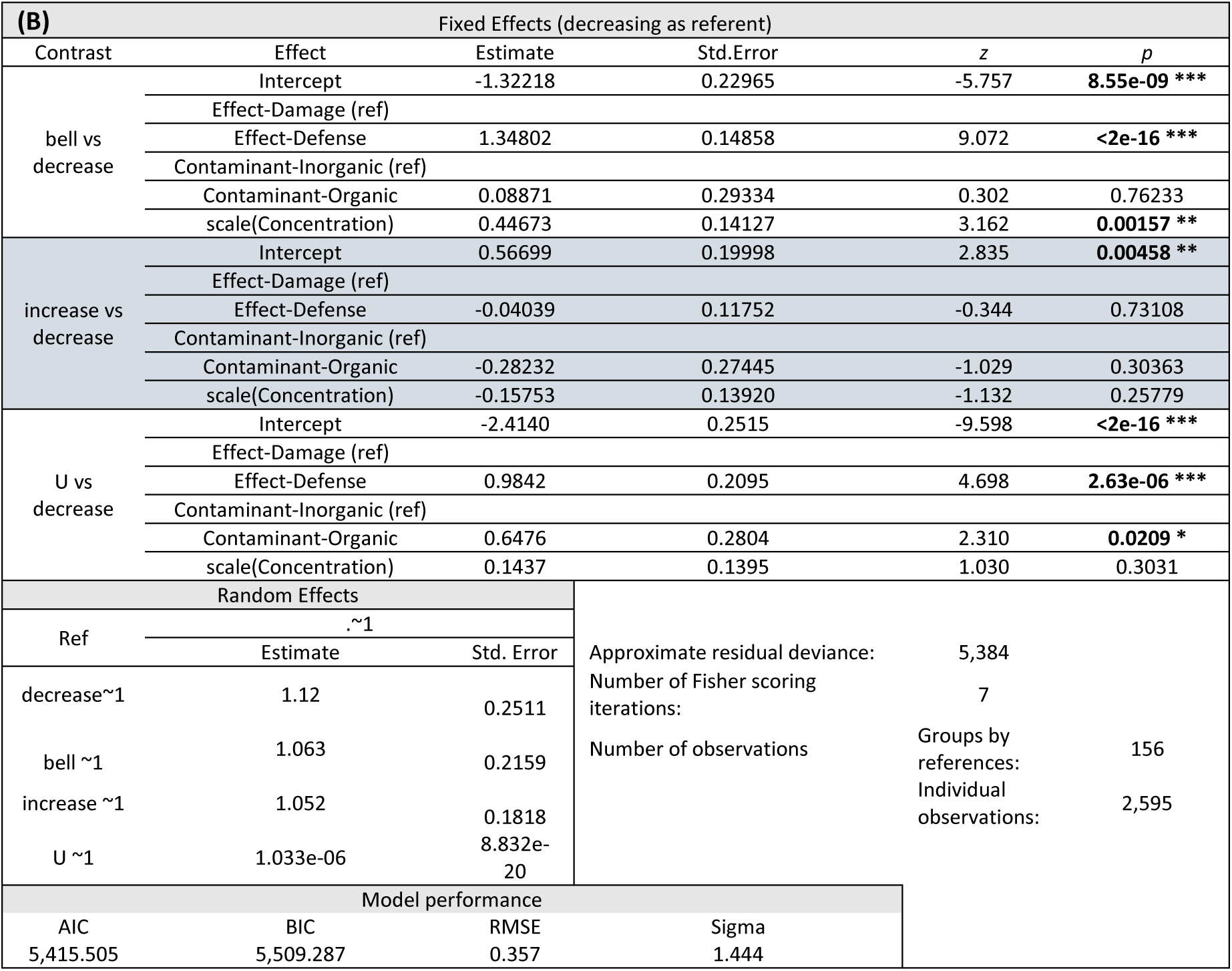
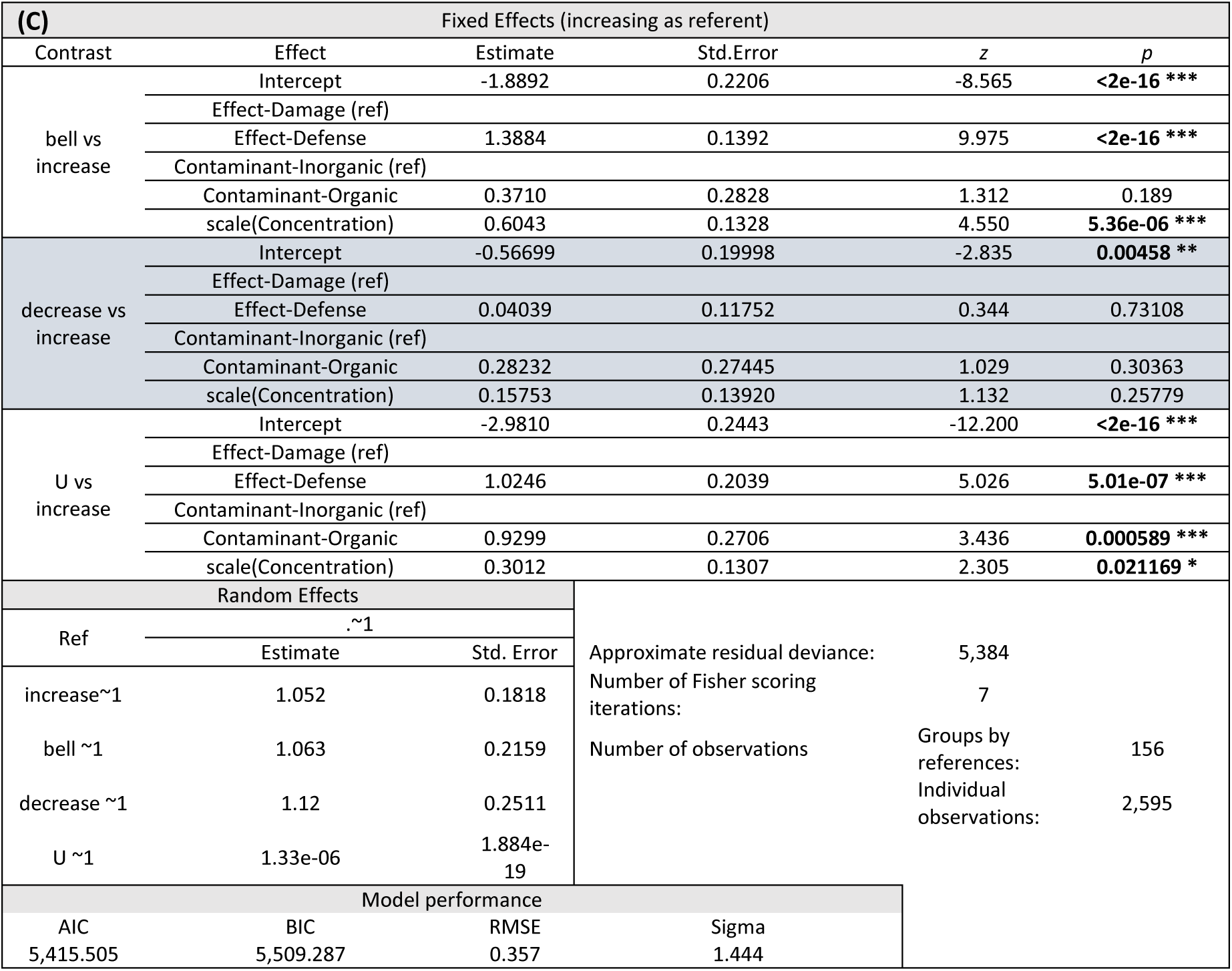
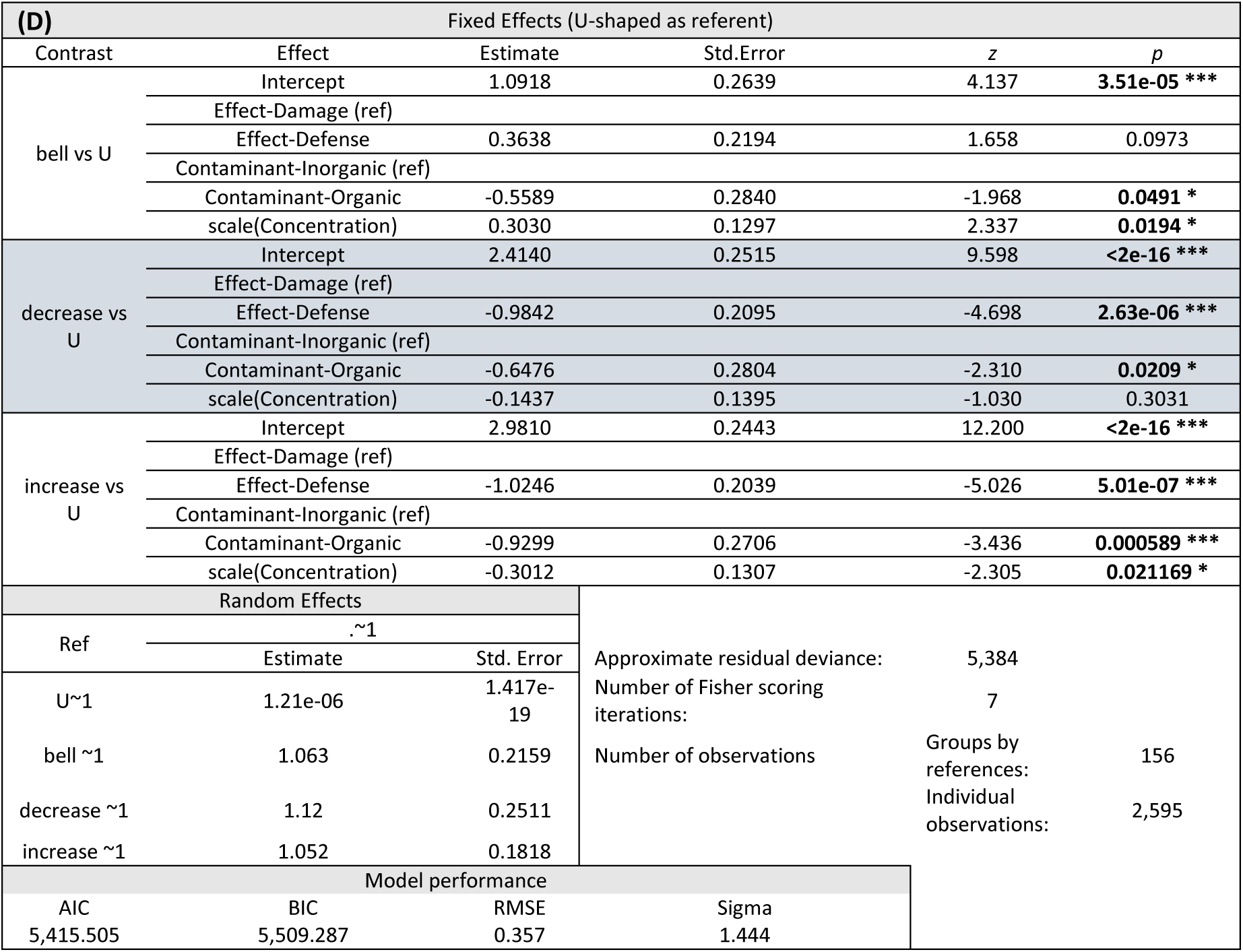
Results of the Multinomial Logistic Regression to examine the effect of the type of biomarkers (Defense vs. Damage), the type of contaminants (Inorganic vs. Organic) and the number of tested concentrations on response trends (bell-shaped, decreasing, increasing and U-shaped).(A) Model with bell-shaped trend as referent, (B) model with decreasing trend as referent, (C) model with increasing trend as referent and (D) model with U-shaped as referent. Significant fixed effects are depicted in bold font and indicated with asterisks as follows: * *p* <0.05, ** *p* < 0.01, *** *p* < 0.001.

## 3.5. Response trends of biomarkers according to the type of contaminant: organic or inorganic

A total of 1,321 observations in the meta-analysis database relates to the effect of organic contaminants and 1,274 to the effect of inorganic contaminants. Biomarkers concerning organic contaminant exposure were either categorized as defense biomarkers (646) or damage biomarkers (675) (Table S1). Among the defense biomarkers, 223 (34.5%) had a bell-shaped trend, 150 (23.2%) had a decreasing trend, 208 (32.26%) had an increasing trend and 65 (10.1%) had a U-shaped trend. The observations were different for damage biomarkers, *i.e.*, 78 (11.6%) had a bell-shaped trend, 224 (33.2%) had a decreasing trend, 342 (50.7%) had an increasing trend and 31 (4.6%) had a U-shaped trend (Fig. 1B). The different biomarkers following inorganic contaminant exposure were distributed with 679 as defense biomarkers and 595 as damage biomarkers (Table S4). Among the defense biomarkers, 179 (26.4 %) had a bell-shaped trend, 149 (21.9%) had a decreasing trend, 321 (47.3%) had an increasing trend and 30 (4.4%) had a U-shaped trend. For damage biomarkers, the observations were different with 614 (10.3%) with a bell-shaped trend, 215 (36.1%) with a decreasing trend, 302 (50.8%) with an increasing trend and 17 (2.9%) with a U-shaped trend (Fig. 1C). The proportion of biomarkers with a biphasic trend (bell-shaped or U-shaped) and the proportion of biomarkers with a monotonic trend (decreasing or increasing) were not affected by the type of contaminant (Tables 2, 3, S3-6 and S8-10). The type of contaminant had only an influence on the proportion of U-shaped trends (Table 3D), with more U-shaped response after an exposure to organic contaminants. In all other cases, the contaminant type had no influence on the type of trends.

### 3.6. Biomarker response trends according to phylum

To determine the effects of taxa on the biomarker response trends, bell- and U-shaped trends were grouped as a biphasic response whereas the increasing and decreasing trends were classified as a monotonic response. These groupings into two main categories were made to have a sufficient number of data for each type of response in each phylum. For the selected phyla, a uniform response is obtained. Defense biomarkers have significantly higher proportions of biphasic trends than damage biomarkers (Fig. 2, and Tables S3-6 and S8-10), except for Cyanobacteria where there is no significant difference among groupings (Table S7). Conversely, damage biomarkers have significantly higher proportions of monotonic trends than defense biomarkers. The type of contaminant (inorganic or organic) had no influence on the response trends.

**Figure 2:**
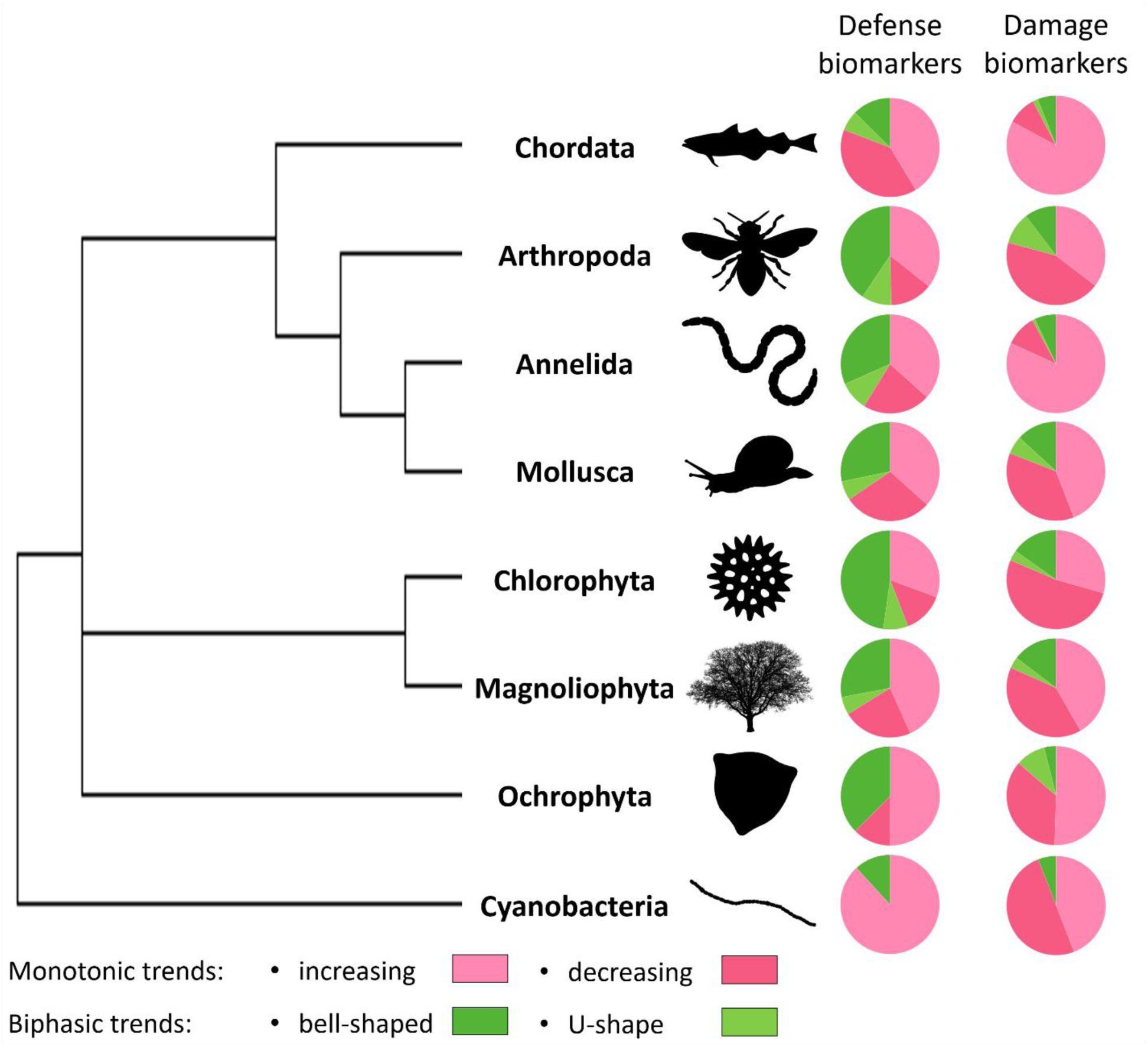
Representation of an unrooted tree of the most represented phyla (with a species example representation) and their proportions of trend of biomarker responses by response mechanism (defense or damage).

### 3.7. Biomarker response trends according to the number of tested concentrations

The number of concentrations used has a significant influence on the observation of biphasic trends. Indeed, the larger it is, the greater the proportion of biphasic trends (bell- and U-shaped) compared to monotonic trends (Tables 2 and 3A-D). Overall, 40% of our observations came from studies that had tested four concentrations, 30% of the observations from five concentrations, 18% from six concentrations and the remainder between 7 and 15 concentrations.

## 4. Discussion

### 4.1. Biomarkers have different response patterns depending on their biological role

The present meta-analysis highlights that defense biomarkers can mainly be described by a bell- or a U-shape dose-response curve, *i.e*., by a biphasic trend, whereas the damage biomarkers are mainly characterized by a linear trend, either increasing or decreasing, *i.e*., by a monotonic trend (Fig. 1A). This difference in response trend could be related to the intrinsic mechanisms of these biomarkers. Molecular, sub-cellular and cellular processes involved in the defense of organisms against stress (here exposure to a contaminant) are induced at low concentrations. Then, the concentrations of defense biomarkers describing a bell-shape curve increase as a function of the increase of contaminant concentration in an attempt to maintain cellular integrity. When the concentrations of the contaminant are too high, these defense mechanisms are overwhelmed and a reduction in their response at the highest concentrations is then often gradually observed[23]. Defense biomarkers with a U-shaped dose-response would follow the same principle of biphasic response but in reverse[23]. The induction of damage biomarkers would be initiated mainly once the defense mechanisms are overwhelmed and would result from the continuous degradation of cellular and sub-cellular compounds, hence their increasing and decreasing trend are proportional to the intensity of the stress[24].

### 4.2. Contaminants trigger similar defense and damage mechanisms in organisms

Trends in defense and damage biomarkers were demonstrated to be independent of the nature of the contaminants (organic or inorganic). Indeed, except for the U-shaped response trends after exposure to an inorganic contaminant (Tables 3A-D), the biomarkers involved in defense mechanisms were mainly described by an increase in bell-shaped and U-shaped trends whereas those involved in damage processes mainly vary in a linear way (Fig. 1B and 1C). This observation supports the tenet that cellular defense mechanisms are not specific to a particular stress[25] but can respond to a multitude of disturbances. In this study, the impact of organic and inorganic contaminants was assessed but similar results might be observed with other stressors such as pH, UV irradiation or temperature. For example, increasing temperature resulted in a GST activity describing a bell-shaped trend in the fish *Sparus aurata*[26] and U-shaped trend in the fresh crab *Aegla longirostri*[27]. In both organisms, CAT activity in their livers and hepatopancreas followed a bell-shaped trend whereas MDA levels in their muscles showed an increasing trend. Cocktail-effects studies also show these patterns of fluctuation specific to defense and damage biomarkers. For example, EROD (CYP450), GST and SOD activities in goldfish exposed to a mixture of norfloxacin and sulfamethoxazole described a bell-shaped response trend with increasing exposure concentrations whereas DNA damage increased strictly monotonically with contaminant concentrations[28].

Here, the parameters considered for each contaminant were the exposure concentrations and the modification of biomarker responses. However, similar trends could also be observed by studying exposure time (for a given concentration). In earthworms (*Eisenia fetida*) exposed to perfluorooctane sulfonamide for 10 days, two defense biomarkers (CYP450 and GST) showed bell-shaped response trends over time[29]. The biomarkers of damage (MDA level and AChE activity) in *Bellamya aeruginosa* exposed for 28 days to different concentrations of Cd and Pb (alone or in a mixture) always varied monotonically (increase) as a function of time whereas defense biomarkers (SOD, CAT, GPx or metallothioneins) described bell- or U-shaped trends over time at certain concentrations[30]. Nevertheless, further analysis of the literature remains to be done to conclude on these trends regarding other stresses, cocktail-effects and exposure time.

### 4.3. Defense and damage mechanisms are preserved throughout life

The specific study of each of the most represented phyla (Annelida, Arthropoda, Chlorophyta, Chordata, Cyanobacteria Magnoliophyta, Mollusca and Ochrophyta) allowed us to observe the same type of response of defense and damage biomarkers. Indeed, the defense biomarkers showed significantly more biphasic responses than the damage biomarkers. Conversely, they showed significantly less monotonic responses (Fig. 2). This was observed for all phyla considered in the analysis (Tables S3-10). As such, Plantae (unicellular and pluricellular) mainly responded to contaminant exposure (organic and inorganic) in a similar way as Animalia. Although the number of observations was lower for Chromista and Bacteria, the responses to exposure to contaminants were also comparable to organisms from the other kingdoms studied. These observations allow to support, that the processes of cellular defense have been conserved during the evolution within different kingdom[31] and that the organizational similarity of cells leads to comparable cell damage between different species.

### 4.4. Importance of a number of tested concentrations in ecotoxicology

This study also demonstrated the importance of having several tested concentrations in ecotoxicology tests. A large range of tested concentrations increases the possibility of observing biphasic trends, potentially leading to a better understanding of the underlying toxicity mechanisms. This is particularly true if the concentration range also includes “low” concentrations[32]. Adding intermediary concentrations with similar concentration intervals would provide a best modelling of the dose-response curves.

### 4.5. Implications for the use of metabolomics in ecotoxicology

Part of the biomarkers compiled in the present meta-analysis are metabolites of low molecular weight. With the increase use of metabolomics and non-targeted approaches in particular, a large number of metabolites as well as their variation can be measured in organisms upon exposure to a contaminant. However, the identification of these metabolites remain challenging as many metabolites remain unannotated in the currently available databases[13]. Assessing their variations via the use of tools such as DRomics for example[17] and analysing their dose-response trends such as performed in the present meta-analysis is an innovative way to evaluate stress in organisms without losing the interest of omics studies and their large number of data[18].

## 5. Conclusion

The present meta-analysis of biomarker response trends showed that the defense mechanisms of living organisms (Animalia, Bacteria, Chromista and Plantae) exposed to different types of contaminants (organic and inorganic) predominantly described biphasic (bell- and U-shaped) dose-responses. In contrast, the damage processes induced by these contaminants were mostly monotonic (increasing and decreasing). The meta-analysis confirms the relevance of dose-response trend analysis as a new omics data processing approach and the identification of CRIDeR and CRIDaR for environmental risk assessment.

## Abbreviations

AIC: Akaike information criterion
BIC: Bayesian information criterion
CAT: catalase
CRIDaR: concentration range inducing damage responses
CRIDeR: concentration range inducing defense responses
HSP: heat-shock protein
LC-MS: liquid chromatography-mass spectrometry
ML: maximum likelihood
N.D.: non-determined
PQL: Penalized Quasi-Likelihood
RMSE: root-mean-square error
SOD: superoxide dismutase.

## Associated content

The detailed dataset used for the meta-analysis, the parameters used to select the random variables in the models and the results of the logistic multinomial regressions on the effects of the type of biomarker, contaminant and number of doses tested on response trends within the different phyla considered, as well as its PRISMA flow diagram are available in the supplementary information section.

## CRediT authorship contribution statement

**Simon Colas**: Conceptualization; Data curation; Formal analysis; Investigation; Methodology; Software; Validation; Visualization; Writing – original draft; Writing – review & editing. **Séverine Le Faucheur**: Conceptualization; Data curation; Funding acquisition; Project administration; Resources; Supervision; Validation; Writing – original draft; Writing – review & editing.

## References

[1] P. Blandin, Bioindicateurs et diagnostic des systèmes écologiques, Bull. Écologie. 17 (1986) 215–307.

[2] C.A.M. Van Gestel, T.C. Van Brummelen, Incorporation of the biomarker concept in ecotoxicology calls for a redefinition of terms, Ecotoxicology. 5 (1996) 217–225. https://doi.org/10.1007/BF00118992.

[3] M. Nordberg, D.M. Templeton, O. Andersen, J.H. Duffus, Glossary of terms used in ecotoxicology (IUPAC Recommendations 2009), Pure Appl. Chem. 81 (2009) 829–970. https://doi.org/10.1515/iupac.81.0008.

[4] M. Zamocky, P.G. Furtmüller, C. Obinger, Evolution of catalases from bacteria to humans, Antioxid. Redox Signal. 10 (2008) 1527–1548. https://doi.org/10.1089/ars.2008.2046.

[5] R.C. Fink, J.G. Scandalios, Molecular evolution and structure–function relationships of the superoxide dismutase gene families in angiosperms and their relationship to other eukaryotic and prokaryotic superoxide dismutases, Arch. Biochem. Biophys. 399 (2002) 19–36. https://doi.org/10.1006/abbi.2001.2739.

[6] R.S. Gupta, B. Singh, Phylogenetic analysis of 70 kD heat shock protein sequences suggests a chimeric origin for the eukaryotic cell nucleus, Curr. Biol. 4 (1994) 1104–1114. https://doi.org/10.1016/S0960-9822(00)00249-9.

[7] A. Dhawan, M. Bajpayee, D. Parmar, Comet assay: a reliable tool for the assessment of DNA damage in different models, Cell Biol. Toxicol. 25 (2009) 5–32. https://doi.org/10.1007/s10565-008-9072-z.

[8] Y. de Lafontaine, F. Gagné, C. Blaise, G. Costan, P. Gagnon, H.M. Chan, Biomarkers in zebra mussels (*Dreissena polymorpha*) for the assessment and monitoring of water quality of the St Lawrence River (Canada), Aquat. Toxicol. 50 (2000) 51–71. https://doi.org/10.1016/S0166-445X(99)00094-6.

[9] J. Artigas, G. Arts, M. Babut, A.B. Caracciolo, S. Charles, A. Chaumot, B. Combourieu, I. Dahllöf, D. Despréaux, B. Ferrari, N. Friberg, J. Garric, O. Geffard, C. Gourlay-Francé, M. Hein, M. Hjorth, M. Krauss, H.J. De Lange, J. Lahr, K.K. Lehtonen, T. Lettieri, M. Liess, S. Lofts, P. Mayer, S. Morin, A. Paschke, C. Svendsen, P. Usseglio-Polatera, N. van den Brink, E. Vindimian, R. Williams, Towards a renewed research agenda in ecotoxicology, Environ. Pollut. 160 (2012) 201–206. https://doi.org/10.1016/j.envpol.2011.08.011.

[10] C. Bedia, Metabolomics in environmental toxicology: Applications and challenges, Trends Environ. Anal. Chem. 34 (2022) e00161. https://doi.org/10.1016/j.teac.2022.e00161.

[11] O. Prat, D. Degli-Esposti, New challenges: Omics technologies in ecotoxicology, in: Ecotoxicology, Elsevier, 2019: pp. 181–208. https://doi.org/10.1016/B978-1-78548-314-1.50006-7.

[12] J.R. Sempionatto, J.A. Lasalde-Ramírez, K. Mahato, J. Wang, W. Gao, Wearable chemical sensors for biomarker discovery in the omics era, Nat. Rev. Chem. 6 (2022) 899–915. https://doi.org/10.1038/s41570-022-00439-w.

[13] D. Dias, O. Jones, D. Beale, B. Boughton, D. Benheim, K. Kouremenos, J.-L. Wolfender, D. Wishart, Current and future perspectives on the structural identification of small molecules in biological systems, Metabolites. 6 (2016) 46. https://doi.org/10.3390/metabo6040046.

[14] T. Cordier, A. Lanzén, L. Apothéloz-Perret-Gentil, T. Stoeck, J. Pawlowski, Embracing environmental genomics and machine learning for routine biomonitoring, Trends Microbiol. 27 (2019) 387–397. https://doi.org/10.1016/j.tim.2018.10.012.

[15] E.J. Dupree, M. Jayathirtha, H. Yorkey, M. Mihasan, B.A. Petre, C.C. Darie, A critical review of bottom-up proteomics: The good, the bad, and the future of this field, Proteomes. 8 (2020) 14. https://doi.org/10.3390/proteomes8030014.

[16] C. Ritz, Toward a unified approach to dose-response modeling in ecotoxicology, Environ. Toxicol. Chem. 29 (2010) 220–229. https://doi.org/10.1002/etc.7.

[17] F. Larras, E. Billoir, V. Baillard, A. Siberchicot, S. Scholz, T. Wubet, M. Tarkka, M. Schmitt-Jansen, M.-L. Delignette-Muller, DRomics: A turnkey tool to support the use of the dose–response framework for omics data in ecological risk assessment, Environ. Sci. Technol. 52 (2018) 14461–14468. https://doi.org/10.1021/acs.est.8b04752.

[18] S. Colas, B. Marie, M. Milhe-Poutingon, M.-C. Lot, A. Boullemant, C. Fortin, S. Le Faucheur, Meta-metabolomic responses of river biofilms to cobalt exposure and use of dose-response model trends as an indicator of effects, BioXriv. (2023). https://doi.org/10.1101/2023.06.19.545533.

[19] R. van der Oost, J. Beyer, N.P.E. Vermeulen, Fish bioaccumulation and biomarkers in environmental risk assessment: a review, Environ. Toxicol. Pharmacol. 13 (2003) 57–149. https://doi.org/10.1016/S1382-6689(02)00126-6.

[20] A. Viarengo, D. Lowe, C. Bolognesi, E. Fabbri, A. Koehler, The use of biomarkers in biomonitoring: A 2-tier approach assessing the level of pollutant-induced stress syndrome in sentinel organisms, Comp. Biochem. Physiol. Part C Toxicol. Pharmacol. 146 (2007) 281300. https://doi.org/10.1016/j.cbpc.2007.04.011.

[21] H. Akaike, Factor analysis and AIC, Psychometrika. 52 (1987) 317–332.

[22] M. Elff, mclogit: Multinomial Logit Models, with or without random effect or overdispersion. R package version 0.9.7, (2021). https://github.com/melff/mclogit/releases/tag/0.9.7.

[23] J.M. Davis, D.J. Svendsgaard, U-Shaped dose-response curves: Their occurrence and implications for risk assessment, J. Toxicol. Environ. Health. 30 (1990) 71–83. https://doi.org/10.1080/15287399009531412.

[24] J.A. Swenberg, E. Fryar-Tita, Y.-C. Jeong, G. Boysen, T. Starr, V.E. Walker, R.J. Albertini, Biomarkers in toxicology and risk assessment: Informing critical dose–response relationships, Chem. Res. Toxicol. 21 (2008) 253–265. https://doi.org/10.1021/tx700408t.

[25] F.L. Mayer, D.J. Versteeg, M.J. McKee, L.C. Folmar, R.L. Graney, D.C. McCume, B.A. Rattner, Physiological and nonspecific biomarkers, in: Biomarkers, CRC Press, 2018: pp. 5– 86.

[26] D. Madeira, C. Vinagre, M.S. Diniz, Are fish in hot water? Effects of warming on oxidative stress metabolism in the commercial species *Sparus aurata*, Ecol. Indic. 63 (2016) 324–331. https://doi.org/10.1016/j.ecolind.2015.12.008.

[27] C. Cerezer, J.W. Leitemperger, A.M.B. do Amaral, B.C. Ferreira, A.T. Marins, V.L. Loro, M.L. Bartholomei-Santos, S. Santos, Raising the water temperature: consequences in behavior and biochemical biomarkers of the freshwater crab *Aegla longirostri* (Crustacea, Anomura), Environ. Sci. Pollut. Res. 27 (2020) 45349–45357. https://doi.org/10.1007/s11356-020-10423-w.

[28] J. Liu, G. Lu, D. Wu, Z. Yan, A multi-biomarker assessment of single and combined effects of norfloxacin and sulfamethoxazole on male goldfish (*Carassius auratus*), Ecotoxicol. Environ. Saf. 102 (2014) 12–17. https://doi.org/10.1016/j.ecoenv.2014.01.014.

[29] S. Zhao, B. Wang, Z. Zhong, T. Liu, T. Liang, J. Zhan, Contributions of enzymes and gut microbes to biotransformation of perfluorooctane sulfonamide in earthworms (*Eisenia fetida*), Chemosphere. 238 (2020) 124619. https://doi.org/10.1016/j.chemosphere.2019.124619.

[30] X. Liu, Q. Chen, N. Ali, J. Zhang, M. Wang, Z. Wang, Single and joint oxidative stress– related toxicity of sediment-associated cadmium and lead on *Bellamya aeruginosa*, Environ. Sci. Pollut. Res. 26 (2019) 24695–24706. https://doi.org/10.1007/s11356-019-05769-9.

[31] T. Nurnberger, F. Brunner, B. Kemmerling, L. Piater, Innate immunity in plants and animals: striking similarities and obvious differences, Immunol. Rev. 198 (2004) 249–266. https://doi.org/10.1111/j.0105-2896.2004.0119.x.

[32] E.J. Calabrese, L.A. Baldwin, The frequency of U-shaped dose responses in the toxicological literature, Toxicol. Sci. 62 (2001) 330–338. https://doi.org/10.1093/toxsci/62.2.330.

[33] J.J. Stegeman, M. Brouwer, T.D.G. Richard, L. Förlin, B.A. Fowler, B.M. Sanders, P.A. van Veld, Molecular responses to environmental contamination: enzyme and protein systems as indicators of chemical exposure and effect, in: Biomark. Biochem. Physiol. Histol. Markers Anthropog. Stress, Lewis Publisher, Boca Raton, FL, USA, 1992: pp. 235–335.

[34] S.G. George, Enzymology and molecular biology of phase II xenobiotic-conjugating enzymes in fish, in: Aquat. Toxicol. Mol. Biochem. Cell. Perspect., Lewis Publisher, 1994: pp. 37–85.

[35] S. Bard, Multixenobiotic resistance as a cellular defense mechanism in aquatic organisms, Aquat. Toxicol. 48 (2000) 357–389. https://doi.org/10.1016/S0166-445X(00)00088-6.

[36] J.L. Hall, Cellular mechanisms for heavy metal detoxification and tolerance, J. Exp. Bot. 53 (2002) 1–11. https://doi.org/10.1093/jexbot/53.366.1.

[37] B.H. Lauterburg, C.V. Smith, H. Hughes, J.R. Mitchell, Determinants of hepatic glutathione turnover: toxicological significance, Trends Pharmacol. Sci. 3 (1982) 245–248.

[38] M.E. Feder, G.E. Hofmann, heat-shock proteins, molecular chaperones, and the stress response: Evolutionary and ecological physiology, Annu. Rev. Physiol. 61 (1999) 243–282. https://doi.org/10.1146/annurev.physiol.61.1.243.

[39] J.F. Payne, A. Mathieu, W. Melvin, L.L. Fancey, Acetylcholinesterase, an old biomarker with a new future? Field trials in association with two urban rivers and a paper mill in Newfoundland, Mar. Pollut. Bull. 32 (1996) 225–231. https://doi.org/10.1016/0025-326X(95)00112-Z.

[40] V. Matozzo, F. Gagné, M.G. Marin, F. Ricciardi, C. Blaise, Vitellogenin as a biomarker of exposure to estrogenic compounds in aquatic invertebrates: A review, Environ. Int. 34 (2008) 531–545. https://doi.org/10.1016/j.envint.2007.09.008.

[41] D.R. Janero, Malondialdehyde and thiobarbituric acid-reactivity as diagnostic indices of lipid peroxidation and peroxidative tissue injury, Free Radic. Biol. Med. 9 (1990) 515–540. https://doi.org/10.1016/0891-5849(90)90131-2.

[42] K.S. Pikula, A.M. Zakharenko, V. Aruoja, K.S. Golokhvast, A.M. Tsatsakis, Oxidative stress and its biomarkers in microalgal ecotoxicology, Curr. Opin. Toxicol. 13 (2019) 8–15. https://doi.org/10.1016/j.cotox.2018.12.006.

[43] D.K. La, J.A. Swenberg, DNA adducts: biological markers of exposure and potential applications to risk assessment, Mutat. Res. Genet. Toxicol. 365 (1996) 129–146. https://doi.org/10.1016/S0165-1110(96)90017-2.

[44] J.A. Heddle, M. Hite, B. Kirkhart, K. Mavournin, J.T. MacGregor, G.W. Newell, M.F. Salamone, The induction of micronuclei as a measure of genotoxicity, Mutat. Res. Genet. Toxicol. 123 (1983) 61–118. https://doi.org/10.1016/0165-1110(83)90047-7.

[45] D.M. Lowe, R.K. Pipe, Contaminant induced lysosomal membrane damage in marine mussel digestive cells: an in vitro study, Aquat. Toxicol. 30 (1994) 357–365.

[46] L. Lagadic, T. Caquet, F. Ramade, The role of biomarkers in environmental assessment (5). Invertebrate populations and communities, Ecotoxicology. 3 (1994) 193–208. https://doi.org/10.1007/BF00117084.

[47] B. Halliwell, J.M. Gutteridge, Free radicals in biology and medicine, USA, 2015.

