## Supplementary Information for "How do biomarkers dance? Specific moves of defense and damage biomarkers for biological interpretation of dose-response model trends"

**TABLES:**

**Table S1**: Data set

**Table S2**: Choice of model and random effects

**Table S3**: Results of the Multinomial Logistic Regression to examine the effect of the type of biomarkers (Defense vs. Damage), the type of contaminants (Inorganic vs. Organic) and the number of tested concentrations on response trends (Biphasic or Monotonic) in Annelida.

**Table S4**: Results of the Multinomial Logistic Regression to examine the effect of the type of biomarkers (Defense vs. Damage), the type of contaminants (Inorganic vs. Organic) and the number of tested concentrations on response trends (Biphasic or Monotonic) in Arthropoda.

**Table S5**: Results of the Multinomial Logistic Regression to examine the effect of the type of biomarkers (Defense vs. Damage), the type of contaminants (Inorganic vs. Organic) and the number of tested concentrations on response trends (Biphasic or Monotonic) in Chlorophyta.

**Table S6**: Results of the Multinomial Logistic Regression to examine the effect of the type of biomarkers (Defense vs. Damage), the type of contaminants (Inorganic vs. Organic) and the number of tested concentrations on response trends (Biphasic or Monotonic) in Chordata.

**Table S7**: Results of the Multinomial Logistic Regression to examine the effect of the type of biomarkers (Defense vs. Damage), the type of contaminants (Inorganic vs. Organic) and the number of tested concentrations on response trends (Biphasic or Monotonic) in Cyanobacteria.

**Table S8**: Results of the Multinomial Logistic Regression to examine the effect of the type of biomarkers (Defense vs. Damage), the type of contaminants (Inorganic vs. Organic) and the number of tested concentrations on response trends (Biphasic or Monotonic) in Magnoliophyta.

**Table S9**: Results of the Multinomial Logistic Regression to examine the effect of the type of biomarkers (Defense vs. Damage), the type of contaminants (Inorganic vs. Organic) and the number of tested concentrations on response trends (Biphasic or Monotonic) in Mollusca.

**Table S10**: Results of the Multinomial Logistic Regression to examine the effect of the type of biomarkers (Defense vs. Damage), the type of contaminants (Inorganic vs. Organic) and the number of tested concentrations on response trends (Biphasic or Monotonic) in Ochrophyta.

**TEXTS:**

**Text S1:** References used for the meta-analysis.

**Text S2:** Abbreviations used in the meta-analysis data set.

**FIGURES:**

**Figure S1**: Meta-analysis PRISMA flow diagram.

**Table S1**: Data set

.xlsx file

The “Biomarker” column corresponds to the known biomarker studied, “Role” to its broad biological role and “Effect” to the type of biomarker (defense or damage), “Organism” to the species studied, “Phylum” and “Kingdom” to its corresponding phylum and kingdom, “Stress” to the contaminant studied, “Contaminant” to the contaminant family (organic or inorganic), “Range” the concentration range tested, “Concentration” the number of concentrations tested, “Exposure time” the exposure time studied, “Trend” the trend of the dose-response curve (bell-shaped, decrease, increase, U-shaped, constant or ND), “Phase” the type of trend (biphasic or monotonic), “Num_ref” the reference number indicated in Text S1 and “Ref” the reference used.”

**Table S2**: Formulas of multinomial logit models with different interactions between fixed and random effect variables and model performance, in bold the models with the best AIC selected for further meta-analysis.

| Equation number | Formula | | Model performance | | | |
|---|---|---|---|---|---|---|
| | Fixed effects | Random effect | AIC | BIC | RMSE | Sigma |
| Biphasic vs. Monotonic | | | | | | |
| 1 | Phase ~ Effect + Contaminant + scale(Concentration) | + ~1\|Ref + ~1\|Time_exposure +~1\|Phylum | 2,488.579 | 2,547.193 | 0.370 | 0.976 |
| 2 | Phase ~ Effect + Contaminant + scale(Concentration) | + ~1\|Time_exposure +~1\|Phylum | 2,651.078 | 2,697.969 | 0.400 | 1.008 |
| 3 | Phase ~ Effect + Contaminant + scale(Concentration) | + ~1\|Ref + ~1\|Time_exposure | 2,486.946 | 2,533.837 | 0.370 | 0.977 |
| 4 | Phase ~ Effect + Contaminant + scale(Concentration) | + ~1\|Ref + ~1\|Phylum | 2,484.674 | 2,531.565 | 0.370 | 0.976 |
| **5** | **Phase ~ Effect + Contaminant + scale(Concentration)** | **+ ~1\|Ref** | 2,483.060 | 2,518.229 | 0.370 | 0.977 |
| 6 | Phase ~ Effect + scale(Concentration) | + ~1\|Ref | 2,483.827 | 2,513.134 | 0.370 | 0.977 |
| 7 | Phase ~ Effect + Contaminant | + ~1\|Ref | 2,499.384 | 2,528.691 | 0.369 | 0.980 |
| 8 | Phase ~ Effect | + ~1\|Ref | 2,499.248 | 2,522.693 | 0.369 | 0.980 |
| Bell-shaped vs. Decreasing / vs. Increasing / vs. U-shaped | | | | | | |
| 9 | Trends ~ Effect + Contaminant + scale(Concentration) | + ~1\|Ref + ~1\|Time_exposure +~1\|Phylum | 5,426.145 | 5,566.818 | 0.357 | 1.443 |
| 10 | Trends ~ Effect + Contaminant + scale(Concentration) | + ~1\|Time_exposure +~1\|Phylum | 5,797.245 | 5,914.472 | 0.384 | 1.493 |
| 11 | Trends ~ Effect + Contaminant + scale(Concentration) | + ~1\|Ref + ~1\|Time_exposure | 5,423.504 | 5,540.731 | 0.357 | 1.444 |
| 12 | Trends ~ Effect + Contaminant + scale(Concentration) | + ~1\|Ref + ~1\|Phylum | 5,418.463 | 5,535.69 | 0.570 | 1.443 |
| **13** | **Trends ~ Effect + Contaminant + scale(Concentration)** | **+ ~1\|Ref** | 5,415.505 | 5,509.287 | 0.357 | 1.444 |
| 14 | Trends ~ Effect + scale(Concentration) | + ~1\|Ref | 5,421.698 | 5,497.895 | 0.357 | 1.444 |
| 15 | Trends ~ Effect + Contaminant | + ~1\|Ref | 5,431.398 | 5,507.596 | 0.357 | 1.446 |
| 16 | Trends ~ Effect | + ~1\|Ref | 5,436.620 | 5,495.233 | 0.570 | 1.446 |

**Table S3**: Results of the Multinomial Logistic Regression to examine the effect of the type of biomarkers (Defense vs. Damage), the type of contaminants (Inorganic vs. Organic) and the number of tested concentrations on response trends (Biphasic or Monotonic) in Annelida. Significant fixed effects are depicted in bold font and indicated with asterisks as follows: * $p < 0.5$, ** $p < 0.01$, *** $p < 0.001$.

| Fixed Effects (Annelida phylum) | | | | | |
|---|---|---|---|---|---|
| Contrast | Effect | Estimate | Std.Error | *z* | *p* |
| monotonic vs biphasic | Intercept | 2.1382 | 0.5674 | 3.769 | **0.000164 ***** |
| | Effect-Damage (ref) | | | | |
| | Effect-Defense | -1.9880 | 0.2942 | -6.758 | **1.4e-11 ***** |
| | Contaminant-Inorganic (ref) | | | | |
| | Contaminant-Organic | 0.3285 | 0.5645 | 0.582 | 0.560638 |
| | scale(Concentration) | -0.2692 | 0.2127 | -1.266 | 0.205674 |

| Random Effects | | | | | |
|---|---|---|---|---|---|
| Ref | .~1 | | Approximate residual deviance: | 531.8 | |
| | Estimate | Std. Error | Number of Fisher scoring iterations: | 6 | |
| biphasic~1 | 0.6125 | 0.06042 | Number of observations | Groups by references: | 25 |
| monotonic ~1 | 0.501 | 0.03056 | | Individual observations: | 552 |

| Model performance | | | |
|---|---|---|---|
| AIC | BIC | RMSE | Sigma |
| 543.779 | 569.660 | 0.377 | 0.985 |

**Table S4**: Results of the Multinomial Logistic Regression to examine the effect of the type of biomarkers (Defense vs. Damage), the type of contaminants (Inorganic vs. Organic) and the number of tested concentrations on response trends (Biphasic or Monotonic) in Arthropoda. Significant fixed effects are depicted in bold font and indicated with asterisks as follows: * $p < 0.5$, ** $p < 0.01$, *** $p < 0.001$.

| Fixed Effects (Arthropoda phylum) | | | | | |
|---|---|---|---|---|---|
| Contrast | Effect | Estimate | Std.Error | *z* | *p* |
| monotonic vs biphasic | Intercept | 1.34861 | 0.45723 | 2.950 | **0.003182 **** |
| | Effect-Damage (ref) | | | | |
| | Effect-Defense | -1.36876 | 0.41989 | -3.260 | **0.001115 **** |
| | Contaminant-Inorganic (ref) | | | | |
| | Contaminant-Organic | 0.03368 | 0.32561 | 0.103 | 0.917626 |
| | scale(Concentration) | -0.54681 | 0.16159 | -3.384 | **0.000715 ***** |

| Random Effects | | | | | |
|---|---|---|---|---|---|
| Ref | .~1 | | Approximate residual deviance: | 235.1 | |
| | Estimate | Std. Error | Number of Fisher scoring iterations: | 4 | |
| biphasic~1 | 3.579e-07 | 1.052e-20 | Number of observations | Groups by references: | 19 |
| monotonic ~1 | 3.701e-07 | 1.163e-20 | | Individual observations: | 191 |

| Model performance | | | |
|---|---|---|---|
| AIC | BIC | RMSE | Sigma |
| 247.129 | 266.643 | 0.464 | 1.121 |

**Table S5**: Results of the Multinomial Logistic Regression to examine the effect of the type of biomarkers (Defense vs. Damage), the type of contaminants (Inorganic vs. Organic) and the number of tested concentrations on response trends (Biphasic or Monotonic) in Chlorophyta. Significant fixed effects are depicted in bold font and indicated with asterisks as follows: * $p$ < 0.5, ** $p$ < 0.01, *** $p$ < 0.001.

| Fixed Effects (Chlorophyta phylum) | | | | | |
|---|---|---|---|---|---|
| Contrast | Effect | Estimate | Std.Error | *z* | *p* |
| monotonic vs biphasic | Intercept | 1.5853 | 0.3725 | 4.256 | **2.08e-05** *** |
| | Effect-Damage (ref) | | | | |
| | Effect-Defense | -1.8137 | 0.3261 | -5.561 | **2.68e-08** *** |
| | Contaminant-Inorganic (ref) | | | | |
| | Contaminant-Organic | -0.4830 | 0.5859 | -0.824 | 0.4097 |
| | scale(Concentration) | -0.4917 | 0.2732 | -1.800 | 0.0719 |

| Random Effects | | | | | |
|---|---|---|---|---|---|
| Ref | .~1 | | Approximate residual deviance: | 334.5 | |
| | Estimate | Std. Error | Number of Fisher scoring iterations: | 6 | |
| biphasic~1 | 0.433 | 0.01933 | Number of observations | Groups by references: | 24 |
| monotonic ~1 | 0.7357 | 0.1192 | | Individual observations: | 332 |

| Model performance | | | |
|---|---|---|---|
| AIC | BIC | RMSE | Sigma |
| 346 | 369.366 | 0.376 | 1.010 |

**Table S6**: Results of the Multinomial Logistic Regression to examine the effect of the type of biomarkers (Defense vs. Damage), the type of contaminants (Inorganic vs. Organic) and the number of tested concentrations on response trends (Biphasic or Monotonic) in Chordata. Significant fixed effects are depicted in bold font and indicated with asterisks as follows: * $p < 0.5$, ** $p < 0.01$, *** $p < 0.001$.

| Fixed Effects (Chordata phylum) | | | | | |
|---|---|---|---|---|---|
| Contrast | Effect | Estimate | Std.Error | *z* | *p* |
| monotonic vs biphasic | Intercept | 3.4602 | 0.6872 | 5.035 | **4.77e-07 *** ** |
| | Effect-Damage (ref) | | | | |
| | Effect-Defense | -1.2579 | 0.5196 | -2.421 | **0.0155 *** |
| | Contaminant-Inorganic (ref) | | | | |
| | Contaminant-Organic | -1.0495 | 0.7317 | -1.434 | 0.1515 |
| | scale(Concentration) | -0.8579 | 0.3361 | -2.553 | **0.0107 *** |
| Random Effects | | | | | |
| Ref | .~1 | | Approximate residual deviance: | 164.4 | |
| | Estimate | Std. Error | Number of Fisher scoring iterations: | 7 | |
| biphasic~1 | 0.2086 | 0.002018 | Number of observations | Groups by references: | 22 |
| monotonic ~1 | 1.201 | 1.235 | | Individual observations: | 283 |
| Model performance | | | | | |
| AIC | BIC | RMSE | Sigma | | |
| 176.384 | 198.256 | 0.268 | 0.768 | | |

**Table S7**: Results of the Multinomial Logistic Regression to examine the effect of the type of biomarkers (Defense vs. Damage), the type of contaminants (Inorganic vs. Organic) and the number of tested concentrations on response trends (Biphasic or Monotonic) in Cyanobacteria. Significant fixed effects are depicted in bold font and indicated with asterisks as follows: * *p* < 0.5, ** *p* < 0.01, *** *p* < 0.001.

| Fixed Effects (Cyanobacteria phylum) | | | | | |
|---|---|---|---|---|---|
| Contrast | Effect | Estimate | Std.Error | *z* | *p* |
| monotonic vs biphasic | Intercept | 3.6065 | 1.1367 | 3.173 | **0.00151 **** |
| | Effect-Damage (ref) | | | | |
| | Effect-Defense | -0.438 | 1.5409 | -0.284 | 0.77609 |
| | Contaminant-Inorganic (ref) | | | | |
| | Contaminant-Organic | | | | |
| | scale(Concentration) | -1.3253 | 0.4180 | -3.170 | **0.00152 **** |

| Random Effects | | |
|---|---|---|
| Ref | .~1 | |
| | Estimate | Std. Error |
| biphasic~1 | 4.723e-08 | 7.45e-23 |
| monotonic ~1 | 4.723e-08 | 7.45e-23 |

| | | |
|---|---|---|
| Approximate residual deviance: | 14.1 | |
| Number of Fisher scoring iterations: | 6 | |
| Number of observations | Groups by references: | 2 |
| | Individual observations: | 51 |

| Model performance | | | |
|---|---|---|---|
| AIC | BIC | RMSE | Sigma |
| 24.098 | 33.757 | 0.180 | 0.542 |

**Table S8**: Results of the Multinomial Logistic Regression to examine the effect of the type of biomarkers (Defense vs. Damage), the type of contaminants (Inorganic vs. Organic) and the number of tested concentrations on response trends (Biphasic or Monotonic) in Magnoliophyta. Significant fixed effects are depicted in bold font and indicated with asterisks as follows: * *p* < 0.5, ** *p* < 0.01, *** *p* < 0.001.

| Fixed Effects (Magnoliophyta phylum) | | | | | |
|---|---|---|---|---|---|
| Contrast | Effect | Estimate | Std.Error | *z* | *p* |
| monotonic vs biphasic | Intercept | 2.2897 | 0.3355 | 6.825 | **8.77e-12 *****|
| | Effect-Damage (ref) | | | | |
| | Effect-Defense | -0.9192 | 0.1994 | -4.610 | **4.03e-06 ***** |
| | Contaminant-Inorganic (ref) | | | | |
| | Contaminant-Organic | -0.9341 | 0.5476 | -1.706 | 0.0881 |
| | scale(Concentration) | -0.4447 | 0.2142 | -2.076 | **0.0379 *** |

| Random Effects | | |
|---|---|---|
| Ref | .~1 | |
| | Estimate | Std. Error |
| biphasic~1 | 0.3854 | 0.01201 |
| monotonic ~1 | 0.6801 | 0.08049 |

| | | | |
|---|---|---|---|
| Approximate residual deviance: | 682.4 | | |
| Number of Fisher scoring iterations: | 7 | | |
| Number of observations | | Groups by references: | 29 |
| | | Individual observations: | 688 |

| Model performance | | | |
|---|---|---|---|
| AIC | BIC | RMSE | Sigma |
| 694.424 | 721.627 | 0.391 | 0.999 |

**Table S9**: Results of the Multinomial Logistic Regression to examine the effect of the type of biomarkers (Defense vs. Damage), the type of contaminants (Inorganic vs. Organic) and the number of tested concentrations on response trends (Biphasic or Monotonic) in Mollusca. Significant fixed effects are depicted in bold font and indicated with asterisks as follows: * *p* < 0.5, ** *p* < 0.01, *** *p* < 0.001.

| Fixed Effects (Mollusca phylum) | | | | | |
|---|---|---|---|---|---|
| Contrast | Effect | Estimate | Std.Error | *z* | *p* |
| monotonic vs biphasic | Intercept | 1.4160 | 0.5532 | 2.560 | **0.0105 *** |
| | Effect-Damage (ref) | | | | |
| | Effect-Defense | -0.8605 | 0.3507 | -2.454 | **0.0141 *** |
| | Contaminant-Inorganic (ref) | | | | |
| | Contaminant-Organic | 0.3316 | 0.7045 | 0.471 | 0.6378 |
| | scale(Concentration) | -0.3543 | 0.3353 | -1.057 | 0.2907 |

| Random Effects | | | | | |
|---|---|---|---|---|---|
| Ref | .~1 | | Approximate residual deviance: | 287.2 | |
| | Estimate | Std. Error | Number of Fisher scoring iterations: | 6 | |
| biphasic~1 | 0.8728 | 0.2621 | Number of observations | Groups by references: | 18 |
| monotonic ~1 | 0.3882 | 0.01566 | | Individual observations: | 278 |

| Model performance | | | |
|---|---|---|---|
| AIC | BIC | RMSE | Sigma |
| 299.184 | 320.95 | 0.386 | 1.024 |

**Table S10**: Results of the Multinomial Logistic Regression to examine the effect of the type of biomarkers (Defense vs. Damage), the type of contaminants (Inorganic vs. Organic) and the number of tested concentrations on response trends (Biphasic or Monotonic) in Ochrophyta. Significant fixed effects are depicted in bold font and indicated with asterisks as follows: * $p < 0.5$, ** $p < 0.01$, *** $p < 0.001$.

| Fixed Effects (Ochrophyta phylum) | | | | | |
|---|---|---|---|---|---|
| Contrast | Effect | Estimate | Std.Error | *z* | *p* |
| monotonic vs biphasic | Intercept | 2.9583 | 0.9794 | 3.020 | **0.00252 ** ** |
| | Effect-Damage (ref) | | | | |
| | Effect-Defense | -2.4108 | 0.8478 | -2.844 | **0.00446 ** ** |
| | Contaminant-Inorganic (ref) | | | | |
| | Contaminant-Organic | -0.6096 | 1.0141 | -0.601 | 0.54777 |
| | scale(Concentration) | -0.4827 | 0.6550 | -0.737 | 0.46122 |
| Random Effects | | | | | |
| Ref | .~1 | | Approximate residual deviance: | 60.26 | |
| | Estimate | Std. Error | Number of Fisher scoring iterations: | 6 | |
| biphasic~1 | 2.499e-07 | 5.515e-21 | Number of observations | Groups by references: | 8 |
| monotonic ~1 | 2.499e-07 | 5.515e-21 | | Individual observations: | 97 |
| Model performance | | | | | |
| AIC | BIC | RMSE | Sigma | | |
| 72.257 | 87.705 | 0.298 | 0.805 | | |

**Text 1**: References used for the meta-analysis.

1. Abassi, S., Wang, H., Ponmani, T. & Ki, J. Small heat shock protein genes of the green algae *Closterium ehrenbergii* : Cloning and differential expression under heat and heavy metal stresses. *Environ. Toxicol.* **34**, 1013–1024 (2019).
2. Aguirre-Martínez, G. V., Buratti, S., Fabbri, E., DelValls, A. T. & Martín-Díaz, M. L. Using lysosomal membrane stability of haemocytes in *Ruditapes philippinarum* as a biomarker of cellular stress to assess contamination by caffeine, ibuprofen, carbamazepine and novobiocin. *J. Environ. Sci.* **25**, 1408–1418 (2013).
3. Ajima, M. N. O., Pandey, P. K., Kumar, K. & Poojary, N. Assessment of mutagenic, hematological and oxidative stress biomarkers in liver of Nile tilapia, *Oreochromis niloticus* (Linnaeus, 1758) in response to sublethal verapamil exposure. *Drug Chem. Toxicol.* **40**, 286–294 (2017).
4. Ajitha, V. *et al.* Effects of zinc and mercury on ROS-mediated oxidative stress-induced physiological impairments and antioxidant responses in the microalga *Chlorella vulgaris*. *Environ. Sci. Pollut. Res.* **28**, 32475–32492 (2021).
5. Akpiri, R. U., Konya, R. S. & Hodges, N. J. Development of cultures of the marine sponge *Hymeniacidon perleve* for genotoxicity assessment using the alkaline comet assay: Metal-induced DNA damage in *Hymeniacidon perleve*. *Environ. Toxicol. Chem.* **36**, 3314–3323 (2017).
6. Al-Fanharawi, A. A., Rabee, A. M. & Al-Mamoori, A. M. J. Biochemical and molecular alterations in freshwater mollusks as biomarkers for petroleum product, domestic heating oil. *Ecotoxicol. Environ. Saf.* **158**, 69–77 (2018).
7. Alho, L. de O. G. *et al.* Photosynthetic, morphological and biochemical biomarkers as tools to investigate copper oxide nanoparticle toxicity to a freshwater chlorophyceae. *Environ. Pollut.* **265**, 114856 (2020).
8. Almeida, A. R. *et al.* Biochemical and behavioral responses of zebrafish embryos to magnetic graphene/nickel nanocomposites. *Ecotoxicol. Environ. Saf.* **186**, 109760 (2019).

9. An, J., Zhou, Q., Sun, F. & Zhang, L. Ecotoxicological effects of paracetamol on seed germination and seedling development of wheat (*Triticum aestivum* L.). *J. Hazard. Mater.* **169**, 751–757 (2009).
10. Arabi, M. & Mahmoodian, F. Comparative toxicity of fresh and expired butachlor to earthworms *Eisenia fetida* in natural soil: Biomarker responses. *Environ. Chem. Ecotoxicol.* **5**, 108–119 (2023).
11. Ashraf, U. & Tang, X. Yield and quality responses, plant metabolism and metal distribution pattern in aromatic rice under lead (Pb) toxicity. *Chemosphere* **176**, 141–155 (2017).
12. Azizullah, A., Richter, P., Jamil, M. & Häder, D.-P. Chronic toxicity of a laundry detergent to the freshwater flagellate *Euglena gracilis*. *Ecotoxicology* **21**, 1957–1964 (2012).
13. Badiou-Bénéteau, A. *et al.* Development of biomarkers of exposure to xenobiotics in the honey bee *Apis mellifera*: Application to the systemic insecticide thiamethoxam. *Ecotoxicol. Environ. Saf.* **82**, 22–31 (2012).
14. Bao, S. *et al.* Effects of diclofenac on the expression of Nrf2 and its downstream target genes in mosquito fish (*Gambusia affinis*). *Aquat. Toxicol.* **188**, 43–53 (2017).
15. Bártová, K., Hilscherová, K., Babica, P., Maršálek, B. & Bláha, L. Effects of microcystin and complex cyanobacterial samples on the growth and oxidative stress parameters in green alga *Pseudokirchneriella subcapitata* and comparison with the model oxidative stressor-herbicide paraquat. *Environ. Toxicol.* **26**, 641–648 (2011).
16. Bertrand, L., Marino, D. J., Monferrán, M. V. & Amé, M. V. Can a low concentration of an organophosphate insecticide cause negative effects on an aquatic macrophyte? Exposure of *Potamogeton pusillus* at environmentally relevant chlorpyrifos concentrations. *Environ. Exp. Bot.* **138**, 139–147 (2017).
17. Brandão, F. P., Pereira, J. L., Gonçalves, F. & Nunes, B. The impact of paracetamol on selected biomarkers of the mollusc species *Corbicula fluminea*: Oxidative Stress in *Corbicula Fluminea*. *Environ. Toxicol.* **29**, 74–83 (2014).

18. Campos, B. G. de *et al.* A preliminary study on multi-level biomarkers response of the tropical oyster *Crassostrea brasiliana* to exposure to the antifouling biocide DCOIT. *Mar. Pollut. Bull.* **174**, 113241 (2022).

19. Canli, E. G. & Canli, M. Antioxidant system biomarkers of freshwater mussel ( *Unio tigridis* ) respond to nanoparticle ($Al_2O_3$ , CuO, $TiO_2$ ) exposures. *Biomarkers* **26**, 434–442 (2021).

20. Canty, M. N., Hutchinson, T. H., Brown, R. J., Jones, M. B. & Jha, A. N. Linking genotoxic responses with cytotoxic and behavioural or physiological consequences: Differential sensitivity of echinoderms (*Asterias rubens*) and marine molluscs (*Mytilus edulis*). *Aquat. Toxicol.* **94**, 68–76 (2009).

21. Cao, X., Song, Y., Kai, J., Yang, X. & Ji, P. Evaluation of EROD and CYP3A4 activities in earthworm *Eisenia fetida* as biomarkers for soil heavy metal contamination. *J. Hazard. Mater.* **243**, 146–151 (2012).

22. Cao, X., Bi, R. & Song, Y. Toxic responses of cytochrome P450 sub-enzyme activities to heavy metals exposure in soil and correlation with their bioaccumulation in *Eisenia fetida*. *Ecotoxicol. Environ. Saf.* **144**, 158–165 (2017).

23. Carneiro, F. E. *et al.* Influence of temperature on the biomarker responses of bullfrog tadpoles (*Lithobates catesbeianus*) to 2-hydroxyatrazine exposure. *Aquat. Toxicol.* **257**, 106468 (2023).

24. Çelekli, A., Kapı, M. & Bozkurt, H. Effect of Cadmium on Biomass, Pigmentation, Malondialdehyde, and Proline of *Scenedesmus quadricauda* var. *longispina*. *Bull. Environ. Contam. Toxicol.* **91**, 571–576 (2013).

25. Çelekli, A., Gültekin, E. & Bozkurt, H. Morphological and Biochemical Responses of *Spirogyra setiformis* Exposed to Cadmium: Water. *CLEAN - Soil Air Water* **44**, 256–262 (2016).

26. Cenkci, S. *et al.* Lead contamination reduces chlorophyll biosynthesis and genomic template stability in *Brassica rapa* L. *Environ. Exp. Bot.* **67**, 467–473 (2010).

27. Chaâbane, M. *et al.* The potential toxic effects of hexavalent chromium on oxidative stress biomarkers and fatty acids profile in soft tissues of *Venus verrucosa*. *Ecotoxicol. Environ. Saf.* **196**, 110562 (2020).

28. Chang, L. W. *et al.* Responses of molecular indicators of exposure in mesocosms: common carp (*Cyprinus carpio*) exposed to the herbicides alachlor and atrazine. *Environ. Toxicol. Chem.* **24**, 190 (2005).

29. Cheng, J., Qiu, H., Chang, Z., Jiang, Z. & Yin, W. The effect of cadmium on the growth and antioxidant response for freshwater algae *Chlorella vulgaris*. *SpringerPlus* **5**, 1290 (2016).

30. Chibee, G. U. *et al.* Effects of cypermethrin as a model chemical on life cycle and biochemical responses of the tropical stingless bee *Meliponula bocandei* Spinola, 1853. *Environ. Adv.* **5**, 100074 (2021).

31. Ciacci, C. *et al.* Effects of sublethal, environmentally relevant concentrations of hexavalent chromium in the gills of *Mytilus galloprovincialis*. *Aquat. Toxicol.* **120–121**, 109–118 (2012).

32. Contardo-Jara, V. & Gessner, M. O. Uptake and physiological effects of the neonicotinoid imidacloprid and its commercial formulation Confidor® in a widespread freshwater oligochaete. *Environ. Pollut.* **264**, 114793 (2020).

33. Cooley, H. Toxicology of dietary uranium in lake whitefish (*Coregonus clupeaformis*). *Aquat. Toxicol.* **48**, 495–515 (2000).

34. Correia, A. T. *et al.* Multi-biomarker approach to assess the acute effects of cerium dioxide nanoparticles in gills, liver and kidney of *Oncorhynchus mykiss*. *Comp. Biochem. Physiol. Part C Toxicol. Pharmacol.* **238**, 108842 (2020).

35. Cruz de Carvalho, R. *et al.* Glyphosate-based herbicide toxicophenomics in marine diatoms: impacts on primary production and physiological fitness. *Appl. Sci.* **10**, 7391 (2020).

36. Cruz de Carvalho, R. *et al.* Effects of Glyphosate-Based Herbicide on Primary Production and Physiological Fitness of the Macroalgae *Ulva lactuca*. *Toxics* **10**, 430 (2022).

37. Coen, W. M. D. & Janssen, C. R. The use of biomarkers in *Daphnia magna* toxicity testing. IV. Cellular Energy Allocation: a new methodology to assess the energy budget of toxicant-stressed *Daphnia* populations.

38. Djebbi, E., Bonnet, D., Pringault, O., Tlili, K. & Yahia, M. N. D. Effects of nickel oxide nanoparticles on survival, reproduction, and oxidative stress biomarkers in the marine calanoid copepod *Centropages ponticus* under short-term exposure. *Environ. Sci. Pollut. Res.* **28**, 21978–21990 (2021).

39. Domingues, I. *et al.* Biomarkers as a tool to assess effects of chromium (VI): Comparison of responses in zebrafish early life stages and adults. *Comp. Biochem. Physiol. Part C Toxicol. Pharmacol.* **152**, 338–345 (2010).

40. Dong, M. *et al.* Toxic effects of 1-decyl-3-methylimidazolium bromide ionic liquid on the antioxidant enzyme system and DNA in zebrafish (*Danio rerio*) livers. *Chemosphere* **91**, 1107–1112 (2013).

41. Duarte, I. A., Pais, M. P., Reis-Santos, P., Cabral, H. N. & Fonseca, V. F. Biomarker and behavioural responses of an estuarine fish following acute exposure to fluoxetine. *Mar. Environ. Res.* **147**, 24–31 (2019).

42. Duarte, B. *et al.* Effects of Propranolol on Growth, Lipids and Energy Metabolism and Oxidative Stress Response of *Phaeodactylum tricornutum*. *Biology* **9**, 478 (2020).

43. Ebenezer, V. & Ki, J.-S. Physiological and biochemical responses of the marine dinoflagellate Prorocentrum minimum exposed to the oxidizing biocide chlorine. *Ecotoxicol. Environ. Saf.* **92**, 129–134 (2013).

44. El Ayeb, N., Béjaoui, M., Muhr, H., Touaylia, S. & Mahmoudi, E. Behaviour and biochemical responses of the marine clam *Ruditapes decussatus* exposed to phosphogypsum. *Environ. Technol.* **42**, 3651–3662 (2021).

45. Elbaz, A., Wei, Y. Y., Meng, Q., Zheng, Q. & Yang, Z. M. Mercury-induced oxidative stress and impact on antioxidant enzymes in *Chlamydomonas reinhardtii*. *Ecotoxicology* **19**, 1285–1293 (2010).

46. Fang, B. *et al.* Toxicity evaluation of 4,4′-di-CDPS and 4,4′-di-CDE on green algae *Scenedesmus obliquus*: growth inhibition, change in pigment content, and oxidative stress. *Environ. Sci. Pollut. Res.* **25**, 15630–15640 (2018).

47. Farmen, E. *et al.* Acute and sub-lethal effects in juvenile Atlantic salmon exposed to low µg/L concentrations of Ag nanoparticles. *Aquat. Toxicol.* **108**, 78–84 (2012).

48. Farooq, M. A. *et al.* Subcellular distribution, modulation of antioxidant and stress-related genes response to arsenic in *Brassica napus* L. *Ecotoxicology* **25**, 350–366 (2016).

49. Feijão, E. *et al.* Fluoxetine arrests growth of the model diatom *Phaeodactylum tricornutum* by increasing oxidative stress and altering energetic and lipid metabolism. *Front. Microbiol.* **11**, 1803 (2020).

50. Gad, M. F., Mossa, A. H., Refaie, A. A., Ibrahim, N. E. & Mohafrash, S. M. M. Benchmark dose and the adverse effects of exposure to pendimethalin at low dose in female rats. *Basic Clin. Pharmacol. Toxicol.* **130**, 301–319 (2022).

51. Gao, M., Qi, Y., Song, W. & Zhou, Q. Biomarker analysis of combined oxytetracycline and zinc pollution in earthworms (*Eisenia fetida*). *Chemosphere* **139**, 229–234 (2015).

52. García-Gómez, C., Obrador, A., González, D., Babín, M. & Fernández, M. D. Comparative study of the phytotoxicity of ZnO nanoparticles and Zn accumulation in nine crops grown in a calcareous soil and an acidic soil. *Sci. Total Environ.* **644**, 770–780 (2018).

53. Geoffroy, L., Dewez, D., Vernet, G. & Popovic, R. Oxyfluorfen toxic effect on *S. obliquus* evaluated by different photosynthetic and enzymatic biomarkers. *Arch. Environ. Contam. Toxicol.* **45**, 445–452 (2003).

54. George, S. G. Cadmium effects on plaice liver xenobiotic and metal detoxication systems: dose-response. *Aquat. Toxicol.* **15**, 303–309 (1989).

55. Gonçalves, S., Almeida, S. F. P., Figueira, E. & Kahlert, M. Valve teratologies and Chl *c* in the freshwater diatom *Tabellaria flocculosa* as biomarkers for metal contamination. *Ecol. Indic.* **101**, 476–485 (2019).

56. Gunderson, M. P., Boyd, H. M., Kelly, C. I., Lete, I. R. & McLaughlin, Q. R. Modulation of endogenous antioxidants by zinc and copper in signal crayfish (*Pacifastacus leniusculus*). *Chemosphere* **275**, 129982 (2021).

57. Gunderson, M. P. *et al.* Response of phase I and II detoxification enzymes, glutathione, metallothionein and acetylcholine esterase to mercury and dimethoate in signal crayfish (*Pacifastacus leniusculus*). *Chemosphere* **208**, 749–756 (2018).

58. Gür, N., Türker, O. C. & Böcük, H. Toxicity assessment of boron (B) by *Lemna minor* L. and *Lemna gibba* L. and their possible use as model plants for ecological risk assessment of aquatic ecosystems with boron pollution. *Chemosphere* **157**, 1–9 (2016).

59. Hazlina, A. Z., Devanthiran, L. & Fatimah, H. Morphological changes and DNA damage in *Chlorella vulgaris* (ULT-M1) induced by $Hg^{2+}$. *Malays. Appl. Biol.* **48**, 27–33 (2019).

60. Herrero, Ó., Aquilino, M., Sánchez-Argüello, P. & Planelló, R. The BPA-substitute bisphenol S alters the transcription of genes related to endocrine, stress response and biotransformation pathways in the aquatic midge *Chironomus riparius* (Diptera, Chironomidae). *PLOS ONE* **13**, e0193387 (2018).

61. Hinojosa-Garro, D., Osten, J. R. & Dzul-Caamal, R. Banded tetra (*Astyanax aeneus*) as bioindicator of trace metals in aquatic ecosystems of the Yucatan Peninsula, Mexico: Experimental biomarkers validation and wild populations biomonitoring. *Ecotoxicol. Environ. Saf.* **195**, 110477 (2020).

62. Hong, Y. *et al.* Evaluation of biomarkers for ecotoxicity assessment by dose-response dynamic models: Effects of nitrofurazone on antioxidant enzymes in the model ciliated protozoan *Euplotes vannus*. *Ecotoxicol. Environ. Saf.* **144**, 552–559 (2017).

63. Hou, J. *et al.* Antioxidant enzyme activities as biomarkers of fluvial biofilm to ZnO NPs ecotoxicity and the Integrated Biomarker Responses (IBR) assessment. *Ecotoxicol. Environ. Saf.* **133**, 10–17 (2016).

64. Hou, K. *et al.* Toxicity evaluation of chlorpyrifos and its main metabolite 3,5,6-trichloro-2-pyridinol (TCP) to *Eisenia fetida* in different soils. *Comp. Biochem. Physiol. Part C Toxicol. Pharmacol.* **259**, 109394 (2022).

65. Jensen, J., Diao, X. & Scott-fordsmand, J. J. Sub-lethal toxicity of the antiparasitic abamectin on earthworms and the application of neutral red retention time as a biomarker. *Chemosphere* **68**, 744–750 (2007).

66. Jia, X., Zhang, H. & Liu, X. Low levels of cadmium exposure induce DNA damage and oxidative stress in the liver of Oujiang colored common carp *Cyprinus carpio* var. *color*. *Fish Physiol. Biochem.* **37**, 97–103 (2011).

67. Jiang, X. *et al.* Ecotoxicity and genotoxicity of polystyrene microplastics on higher plant Vicia faba. *Environ. Pollut.* **250**, 831–838 (2019).

68. Jiang, H. *et al.* Biphasic dose–response of components from *Coptis chinensis* on feeding and detoxification enzymes of *Spodoptera litura* larvae. *Dose-Response* **18**, 155932582091634 (2020).

69. Jmii, S. & Dewez, D. Toxic Responses of Palladium Accumulation in Duckweed ( *Lemna minor* ): Determination of Biomarkers. *Environ. Toxicol. Chem.* **40**, 1630–1638 (2021).

70. Kanoun-Boulé, M., Vicente, J. A. F., Nabais, C., Prasad, M. N. V. & Freitas, H. Ecophysiological tolerance of duckweeds exposed to copper. *Aquat. Toxicol.* **91**, 1–9 (2009).

71. Kim, B.-M. *et al.* Heavy metals induce oxidative stress and trigger oxidative stress-mediated heat shock protein (hsp) modulation in the intertidal copepod *Tigriopus japonicus*. *Comp. Biochem. Physiol. Part C Toxicol. Pharmacol.* **166**, 65–74 (2014).

72. Kim, W.-K. *et al.* Integration of multi-level biomarker responses to cadmium and benzo[k]fluoranthene in the pale chub (*Zacco platypus*). *Ecotoxicol. Environ. Saf.* **110**, 121–128 (2014).

73. Kirici, M. Toxic effects of copper sulphate pentahydrate on antioxidant enzyme activities and lipid peroxidation of freshwater fish *Capoeta umbla* (Heckel, 1843) tissues. *Appl. Ecol. Environ. Res.* **15**, 1685–1696 (2017).

74. Koppel, D. J., Adams, M. S., King, C. K. & Jolley, D. F. Chronic toxicity of an environmentally relevant and equitoxic ratio of five metals to two Antarctic marine microalgae shows complex mixture interactivity. *Environ. Pollut.* **242**, 1319–1330 (2018).

75. Körpe, D. A. & Aras, S. Evaluation of copper-induced stress on eggplant (*Solanum melongena* L.) seedlings at the molecular and population levels by use of various biomarkers. *Mutat. Res. Toxicol. Environ. Mutagen.* **719**, 29–34 (2011).

76. Kovačević, M., Hackenberger, D. K., Lončarić, Ž. & Hackenberger, B. K. Measurement of multixenobiotic resistance activity in enchytraeids as a tool in soil ecotoxicology. *Chemosphere* **279**, 130549 (2021).

77. Li, M. *et al.* Copper and zinc induction of lipid peroxidation and effects on antioxidant enzyme activities in the microalga *Pavlova viridis* (Prymnesiophyceae). *Chemosphere* **62**, 565–572 (2006).

78. Li, M. *et al.* Cobalt and manganese stress in the microalga *Pavlova viridis* (Prymnesiophyceae): Effects on lipid peroxidation and antioxidant enzymes. *J. Environ. Sci.* **19**, 1330–1335 (2007).

79. Li, H. *et al.* Effects of waterborne nano-iron on medaka (*Oryzias latipes*): Antioxidant enzymatic activity, lipid peroxidation and histopathology. *Ecotoxicol. Environ. Saf.* **72**, 684–692 (2009).

80. Li, X., Yang, Y., Jia, L., Chen, H. & Wei, X. Zinc-induced oxidative damage, antioxidant enzyme response and proline metabolism in roots and leaves of wheat plants. *Ecotoxicol. Environ. Saf.* **89**, 150–157 (2013).

81. Li, X. *et al.* Mesotrione-induced oxidative stress and DNA damage in earthworms (*Eisenia fetida*). *Ecol. Indic.* **95**, 436–443 (2018).

82. Li, X., Hua, Z., Zhang, J. & Gu, L. Ecotoxicological responses and removal of submerged macrophyte *Hydrilla verticillate* to multiple perfluoroalkyl acid (PFAA) pollutants in aquatic environments. *Sci. Total Environ.* **825**, 153919 (2022).

83. Li, P., Zhang, J., Sun, X., Agathokleous, E. & Zheng, G. Atmospheric Pb induced hormesis in the accumulator plant *Tillandsia usneoides*. *Sci. Total Environ.* **811**, 152384 (2022).

84. Liu, Y., Zhou, Q., Xie, X., Lin, D. & Dong, L. Oxidative stress and DNA damage in the earthworm *Eisenia fetida* induced by toluene, ethylbenzene and xylene. *Ecotoxicology* **19**, 1551–1559 (2010).

85. Liu, J., Lu, G., Wu, D. & Yan, Z. A multi-biomarker assessment of single and combined effects of norfloxacin and sulfamethoxazole on male goldfish (*Carassius auratus*). *Ecotoxicol. Environ. Saf.* **102**, 12–17 (2014).

86. Liu, S., Ding, R. & Nie, X. Assessment of oxidative stress of paracetamol to *Daphnia magna* via determination of Nrf1 and genes related to antioxidant system. *Aquat. Toxicol.* **211**, 73–80 (2019).

87. Liu, F., Lu, Z., Wu, H. & Ji, C. Dose-dependent effects induced by cadmium in polychaete *Perinereis aibuhitensis*. *Ecotoxicol. Environ. Saf.* **169**, 714–721 (2019).

88. Liu, X. *et al.* Single and joint oxidative stress–related toxicity of sediment-associated cadmium and lead on *Bellamya aeruginosa*. *Environ. Sci. Pollut. Res.* **26**, 24695–24706 (2019).

89. Lu, G. H., Wang, C. & Zhu, Z. The dose–response relationships for EROD and GST induced by polyaromatic hydrocarbons in *Carassius auratus*. *Bull. Environ. Contam. Toxicol.* **82**, 194–199 (2009).

90. Lüchmann, K. H. *et al.* Biochemical biomarkers and hydrocarbons concentrations in the mangrove oyster *Crassostrea brasiliana* following exposure to diesel fuel water-accommodated fraction. *Aquat. Toxicol.* **105**, 652–660 (2011).

91. Malec, P., Maleva, M. G., Prasad, M. N. V. & Strzałka, K. Responses of <i>Lemna trisulca<i> L. (Duckweed) exposed to low doses of cadmium: thiols, metal binding complexes, and photosynthetic pigments as sensitive biomarkers of ecotoxicity. *Protoplasma* **240**, 69–74 (2010).

92. Martinez, R. S., Sáenz, M. E., Alberdi, J. L. & Di Marzio, W. D. Comparative ecotoxicity of single and binary mixtures exposures of nickel and zinc on growth and biomarkers of *Lemna gibba*. *Ecotoxicology* **28**, 686–697 (2019).

93. Melegari, S. P., Perreault, F., Costa, R. H. R., Popovic, R. & Matias, W. G. Evaluation of toxicity and oxidative stress induced by copper oxide nanoparticles in the green alga *Chlamydomonas reinhardtii*. *Aquat. Toxicol.* **142–143**, 431–440 (2013).

94. Morais, L. G. *et al.* Multilevel assessment of chlorothalonil sediment toxicity to Latin American estuarine biota: Effects on biomarkers, reproduction and survival in different benthic organisms. *Sci. Total Environ.* **872**, 162215 (2023).

95. Mukherjee, A. *et al.* Tolerance of arsenate-induced stress in *Aspergillus niger*, a possible candidate for bioremediation. *Ecotoxicol. Environ. Saf.* **73**, 172–182 (2010).

96. Navarrete, A. *et al.* Copper excess detoxification is mediated by a coordinated and complementary induction of glutathione, phytochelatins and metallothioneins in the green seaweed *Ulva compressa*. *Plant Physiol. Biochem.* **135**, 423–431 (2019).

97. Nie, X., Wang, X., Chen, J., Zitko, V. & An, T. Response of the freshwater alga *Chlorella vulgaris* to trichloroisocyanuric acid and ciprofloxacin. *Environ. Toxicol. Chem.* **27**, 168 (2008).

98. Oliveira, M., Gravato, C. & Guilhermino, L. Acute toxic effects of pyrene on *Pomatoschistus microps* (Teleostei, Gobiidae): Mortality, biomarkers and swimming performance. *Ecol. Indic.* **19**, 206–214 (2012).

99. Osterauer, R., Faßbender, C., Braunbeck, T. & Köhler, H.-R. Genotoxicity of platinum in embryos of zebrafish (*Danio rerio*) and ramshorn snail (Marisa cornuarietis). *Sci. Total Environ.* **409**, 2114–2119 (2011).

100. Patel, A., Tiwari, S. & Prasad, S. M. Toxicity assessment of arsenate and arsenite on growth, chlorophyll a fluorescence and antioxidant machinery in *Nostoc muscorum*. *Ecotoxicol. Environ. Saf.* **157**, 369–379 (2018).

101. Qian, J. *et al.* Phytotoxicity and oxidative stress of perfluorooctanesulfonate to two riparian plants: *Acorus calamus* and *Phragmites communis*. *Ecotoxicol. Environ. Saf.* **180**, 215–226 (2019).

102. Rai, U. N., Singh, N. K., Upadhyay, A. K. & Verma, S. Chromate tolerance and accumulation in *Chlorella vulgaris* L.: Role of antioxidant enzymes and biochemical changes in detoxification of metals. *Bioresour. Technol.* **136**, 604–609 (2013).

103. Ramadass, K., Megharaj, M., Venkateswarlu, K. & Naidu, R. Toxicity of diesel water accommodated fraction toward microalgae, *Pseudokirchneriella subcapitata* and *Chlorella* sp. MM3. *Ecotoxicol. Environ. Saf.* **142**, 538–543 (2017).

104. Ran, X. *et al.* Assessment of growth rate, chlorophyll *a* fluorescence, lipid peroxidation and antioxidant enzyme activity in *Aphanizomenon flos-aquae*, *Pediastrum simplex* and *Synedra acus* exposed to cadmium. *Ecotoxicology* **24**, 468–477 (2015).

105. Razinger, J., Dermastia, M., Koce, J. D. & Zrimec, A. Oxidative stress in duckweed (*Lemna minor* L.) caused by short-term cadmium exposure. *Environ. Pollut.* **153**, 687–694 (2008).

106. Reinecke, S. A. & Reinecke, A. J. The comet assay as biomarker of heavy metal genotoxicity in earthworms. *Arch. Environ. Contam. Toxicol.* **46**, 208–215 (2004).

107. Rijstenbil, J. Interactions of algal ligands, metal complexation and availability, and cell responses of the diatom *Ditylum brightwellii* with a gradual increase in copper. *Aquat. Toxicol.* **56**, 115–131 (2002).

108. Romero, N. *et al.* Physiological and morphological responses of green microalgae *Chlorella vulgaris* to silver nanoparticles. *Environ. Res.* **189**, 109857 (2020).

109. Sáenz, M. E., Marzio, W. D. D. & Alberdi, J. L. Assessment of Cyfluthrin commercial formulation on growth, photosynthesis and catalase activity of green algae. *Pestic. Biochem. Physiol.* **104**, 50–57 (2012).

110. Sathasivam, R., Ebenezer, V., Guo, R. & Ki, J.-S. Physiological and biochemical responses of the freshwater green algae *Closterium ehrenbergii* to the common disinfectant chlorine. *Ecotoxicol. Environ. Saf.* **133**, 501–508 (2016).

111. Sellami, B. *et al.* Effects of 2-(4-Methoxyphenyl)-5, 6-trimethylene-4H-1, 3, 2-oxathiaphosphorine-2-sulfide on biomarkers of Mediterranean clams *Ruditapes decussatus*. *Ecotoxicol. Environ. Saf.* **120**, 263–269 (2015).

112. Sendra, M., Yeste, M. P., Gatica, J. M., Moreno-Garrido, I. & Blasco, J. Direct and indirect effects of silver nanoparticles on freshwater and marine microalgae (*Chlamydomonas reinhardtii* and *Phaeodactylum tricornutum*). *Chemosphere* **179**, 279–289 (2017).

113. Seth, C. S., Misra, V., Chauhan, L. K. S. & Singh, R. R. Genotoxicity of cadmium on root meristem cells of *Allium cepa*: cytogenetic and Comet assay approach. *Ecotoxicol. Environ. Saf.* **71**, 711–716 (2008).

114. Shi, Z. *et al.* Effects of biochar and thermally treated biochar on *Eisenia fetida* survival, growth, lysosomal membrane stability and oxidative stress. *Sci. Total Environ.* **770**, 144778 (2021).

115. Simon, D. F., Descombes, P., Zerges, W. & Wilkinson, K. J. Global expression profiling of *Chlamydomonas reinhardtii* exposed to trace levels of free cadmium. *Environ. Toxicol. Chem.* **27**, 1668 (2008).

116. Simon, R. *et al.* Mass spectrometry assay as an alternative to the enzyme-linked immunosorbent assay test for biomarker quantitation in ecotoxicology: Application to vitellogenin in Crustacea (*Gammarus fossarum*). *J. Chromatogr. A* **1217**, 5109–5115 (2010).

117. Singh, V., Pandey, B. & Suthar, S. Phytotoxicity of amoxicillin to the duckweed *Spirodela polyrhiza*: Growth, oxidative stress, biochemical traits and antibiotic degradation. *Chemosphere* **201**, 492–502 (2018).

118. Song, P. *et al.* Phthalate induced oxidative stress and DNA damage in earthworms (*Eisenia fetida*). *Environ. Int.* **129**, 10–17 (2019).

119. Stefano, B., Ilaria, C. & Silvano, F. Cholinesterase activities in the scallop *Pecten jacobaeus*: Characterization and effects of exposure to aquatic contaminants. *Sci. Total Environ.* **392**, 99–109 (2008).

120. Stoiber, T. L., Shafer, M. M., Perkins, D. A. K., Hemming, J. D. C. & Armstrong, D. E. Analysis of glutathione endpoints for measuring copper stress in *Chlamydomonas reinhardtii*. *Environ. Toxicol. Chem.* **26**, 1563 (2007).

121. Touaylia, S., Ali, M., Abdellhafidh, K. & Bejaoui, M. Permethrin induced oxidative stress and neurotoxicity on the freshwater beetle *Laccophilus minutus*. *Chem. Ecol.* **35**, 459–471 (2019).

122. Tu, H. T. *et al.* Acetylcholinesterase activity as a biomarker of exposure to antibiotics and pesticides in the black tiger shrimp (*Penaeus monodon*). *Ecotoxicol. Environ. Saf.* **72**, 1463–1470 (2009).

123. Venn, A. A., Quinn, J., Jones, R. & Bodnar, A. P-glycoprotein (multi-xenobiotic resistance) and heat shock protein gene expression in the reef coral *Montastraea franksi* in response to environmental toxicants. *Aquat. Toxicol.* **93**, 188–195 (2009).

124. Vidal, M.-L., Bassères, A. & Narbonne, J.-F. Potential biomarkers of trichloroethylene and toluene exposure in *Corbicula fluminea*. *Environ. Toxicol. Pharmacol.* **9**, 87–97 (2001).

125. Wang, M. & Wang, G. Oxidative damage effects in the copepod *Tigriopus japonicus* Mori experimentally exposed to nickel. *Ecotoxicology* **19**, 273–284 (2010).

126. Wang, C. *et al.* Lead-contaminated soil induced oxidative stress, defense response and its indicative biomarkers in roots of *Vicia faba* seedlings. *Ecotoxicology* **19**, 1130–1139 (2010).

127. Wang, C. *et al.* Stress response and potential biomarkers in spinach (*Spinacia oleracea* L.) seedlings exposed to soil lead. *Ecotoxicol. Environ. Saf.* **74**, 41–47 (2011).

128. Wang, C. *et al.* Detoxification mechanisms, defense responses, and toxicity threshold in the earthworm *Eisenia foetida* exposed to ciprofloxacin-polluted soils. *Sci. Total Environ.* **612**, 442–449 (2018).

129. Wang, X. *et al.* Multi-level ecotoxicological effects of imidacloprid on earthworm (*Eisenia fetida*). *Chemosphere* **219**, 923–932 (2019).

130. Wang, H. *et al.* Biochemical responses and DNA damage induced by herbicide QYR301 in earthworm (*Eisenia fetida*). *Chemosphere* **244**, 125512 (2020).

131. Wang, Z. *et al.* Antioxidant defense system responses, lysosomal membrane stability and DNA damage in earthworms ( *Eisenia fetida* ) exposed to perfluorooctanoic acid: an integrated biomarker approach to evaluating toxicity. *RSC Adv.* **11**, 26481–26492 (2021).

132. Wei, F., Su, T., Wang, D., Li, H. & You, J. Transcriptomic analysis reveals common pathways and biomarkers associated with oxidative damage caused by mitochondrial toxicants in *Chironomus dilutus*. *Chemosphere* **254**, 126746 (2020).

133. Wei, C. *et al.* Hormetic effects of zinc on growth and antioxidant defense system of wheat plants. *Sci. Total Environ.* **807**, 150992 (2022).

134. Xiao, A. *et al.* Carbon and Metal Quantum Dots toxicity on the microalgae *Chlorella pyrenoidosa*. *Ecotoxicol. Environ. Saf.* **133**, 211–217 (2016).

135. Xiong, Q. *et al.* Sublethal or not? Responses of multiple biomarkers in *Daphnia magna* to single and joint effects of BDE-47 and BDE-209. *Ecotoxicol. Environ. Saf.* **164**, 164–171 (2018).

136. Xu, X. *et al.* Toxicological effects, mechanisms, and implied toxicity thresholds in the roots of *Vicia faba* L. seedlings grown in copper-contaminated soil. *Environ. Sci. Pollut. Res.* **22**, 13858–13869 (2015).

137. Xu, Q. *et al.* Response of *Spirodela polyrhiza* to cerium: subcellular distribution, growth and biochemical changes. *Ecotoxicol. Environ. Saf.* **139**, 56–64 (2017).

138. Xu, Z., Yang, Z., Shu, W. & Zhu, T. Combined toxicity of soil antimony and cadmium on earthworm *Eisenia fetida*: Accumulation, biomarker responses and joint effect. *J. Hazard. Mater. Lett.* **2**, 100018 (2021).

139. Yang, J., Cao, J., Xing, G. & Yuan, H. Lipid production combined with biosorption and bioaccumulation of cadmium, copper, manganese and zinc by oleaginous microalgae Chlorella minutissima UTEX2341. *Bioresour. Technol.* **175**, 537–544 (2015).

140. Yang, W. *et al.* Chemical- and species-specific toxicity of nonylphenol and octylphenol to microalgae *Chlorella pyrenoidosa* and *Scenedesmus obliquus*. *Environ. Toxicol. Pharmacol.* **81**, 103517 (2021).

141. Yang, H. *et al.* Ecotoxicological and biochemical effects of di(2-ethylhexyl)phthalate on wheat (Jimai 22, *Triticum aestivum* L.). *J. Hazard. Mater.* **447**, 130816 (2023).

142. Yao, X. *et al.* Extreme environmental doses of diisobutyl phthalate exposure induce oxidative stress and DNA damage in earthworms (*Eisenia fetida*): Evidence at the biochemical and molecular levels. *J. Environ. Manage.* **331**, 117321 (2023).

143. Yu, H., Zhang, X., Hu, J., Peng, J. & Qu, J. Ecotoxicity of polystyrene microplastics to submerged carnivorous *Utricularia vulgaris* plants in freshwater ecosystems. *Environ. Pollut.* **265**, 114830 (2020).

144. Zhan, J., Wang, S., Li, F., Ji, C. & Wu, H. Dose-dependent responses of metabolism and tissue injuries in clam *Ruditapes philippinarum* after subchronic exposure to cadmium. *Sci. Total Environ.* **779**, 146479 (2021).

145. Zhang, W. *et al.* Earthworm cytochrome P450 determination and application as a biomarker for diagnosing PAH exposure. *J. Environ. Monit.* **8**, 963 (2006).

146. Zhang, Y. *et al.* Long-term toxicity effects of cadmium and lead on *Bufo raddei* tadpoles. *Bull. Environ. Contam. Toxicol.* **79**, 178–183 (2007).

147. Zhang, S., Zhang, H., Qin, R., Jiang, W. & Liu, D. Cadmium induction of lipid peroxidation and effects on root tip cells and antioxidant enzyme activities in *Vicia faba* L. *Ecotoxicology* **18**, 814–823 (2009).

148. Zhang, T. *et al.* Mercury induced oxidative stress, DNA damage, and activation of antioxidative system and Hsp70 induction in duckweed ( *Lemna minor* ). *Ecotoxicol. Environ. Saf.* **143**, 46–56 (2017).

149. Zhang, Y. *et al.* Cadmium-induced oxidative stress, histopathology, and transcriptome changes in the hepatopancreas of freshwater crayfish (*Procambarus clarkii*). *Sci. Total Environ.* **666**, 944–955 (2019).

150. Zhang, H. *et al.* Benzotriazole alleviates copper mediated lysosomal membrane damage and antioxidant defense system responses in earthworms (*Eisenia fetida*). *Ecotoxicol. Environ. Saf.* **197**, 110618 (2020).

151. Zhang, C. *et al.* Applying fungicide on earthworms: Biochemical effects of *Eisenia fetida* exposed to fluoxastrobin in three natural soils. *Environ. Pollut.* **258**, 113666 (2020).

152. Zhang, M., Chen, J., Li, Y., Li, G. & Zhang, Z. Sub-chronic ecotoxicity of triphenyl phosphate to earthworms (*Eisenia fetida*) in artificial soil: Oxidative stress and DNA damage. *Ecotoxicol. Environ. Saf.* **241**, 113796 (2022).

153. Zhao, S., Wang, Y. & Duo, L. Biochemical toxicity, lysosomal membrane stability and DNA damage induced by graphene oxide in earthworms. *Environ. Pollut.* **269**, 116225 (2021).

154. Zheng, S., Wang, Y., Zhou, Q. & Chen, C. Responses of Oxidative Stress Biomarkers and DNA Damage on a Freshwater Snail (*Bellamya aeruginosa*) Stressed by Ethylbenzene. *Arch. Environ. Contam. Toxicol.* **65**, 251–259 (2013).

155. Zheng, Y., Zhou, K., Tang, J., Liu, C. & Bai, J. Impacts of di-(2-ethylhexyl) phthalate on *Folsomia candida* (Collembola) assessed with a multi-biomarker approach. *Ecotoxicol. Environ. Saf.* **232**, 113251 (2022).

156. Zhou, Z. *et al.* Glutathione S-transferase (GST) genes from marine copepods *Acartia tonsa*: cDNA cloning and mRNA expression in response to 1,2-dimethylnaphthalene. *Aquat. Toxicol.* **224**, 105480 (2020).

**Text 2**: Abbreviations used in the meta-analysis data set.

5-HTR, 5-hydroxytryptamine receptor; 8-OHdG, 8-hydroxydesoxyguanosine; ABS/RC, absorption per active reaction center; AChE, acetylcholinesterase; ALP, alkaline phosphatase; ALT, alanine transaminase; AMP, adenosine monophosphate; AOEC, active oxygen-evolving complex; APx, ascorbate peroxidase; AsA, ascorbic acid; AST, aspartate transaminase; ATP, adenosine triphosphate; CaE, carboxylesterase; CAT, catalase; CEA, cellular energy allocation; ChE, cholinesterase; COI, cytochrome oxidase subunit I; CYP, cytochrome; DHA, docosahexaenoic acid; DW, dry weight; EP, endoprotease; EROD, 7-ethoxyresorufin-O-deethylation; FW, fresh weight; G6PDH, glucose-6-phosphate dehydrogenase; GCLC, glutamate-cysteine ligase catalytic subunit; GGT, gamma-glutamyl transferase; GK, glucokinase; GPx, glutathione peroxidase; GR, glutathione reductase; GSH, glutathione; GSSH, glutathione disulfide; GST, glutathione-S-transferase; HSP, heat-shock protein; LDH, lactate dehydrogenase; LOx, lipoxygenase; LPO, lipid peroxidation; MDA, malondialdehyde; MROD, 7-methoxyresofurin-O-demethylation; MT, metallothionein; MXR, multi-xenobiotic resistance; NADH, reduced nicotinamide adenine dinucleotide; NP, nanoparticle; NRRT, neutral red retention time; OTM, olive tail moment; PDH, pyruvate dehydrogenase; POD, peroxidase; PS, photosystem; ROS, reactive oxygen species; T-AOC, total antioxidant capacity; TBARS, thiobarbituric acid reactive substance; UGP, UDP-glucuronosyltransferase.

**Figure S1:** Meta-analysis PRISMA flow diagram.

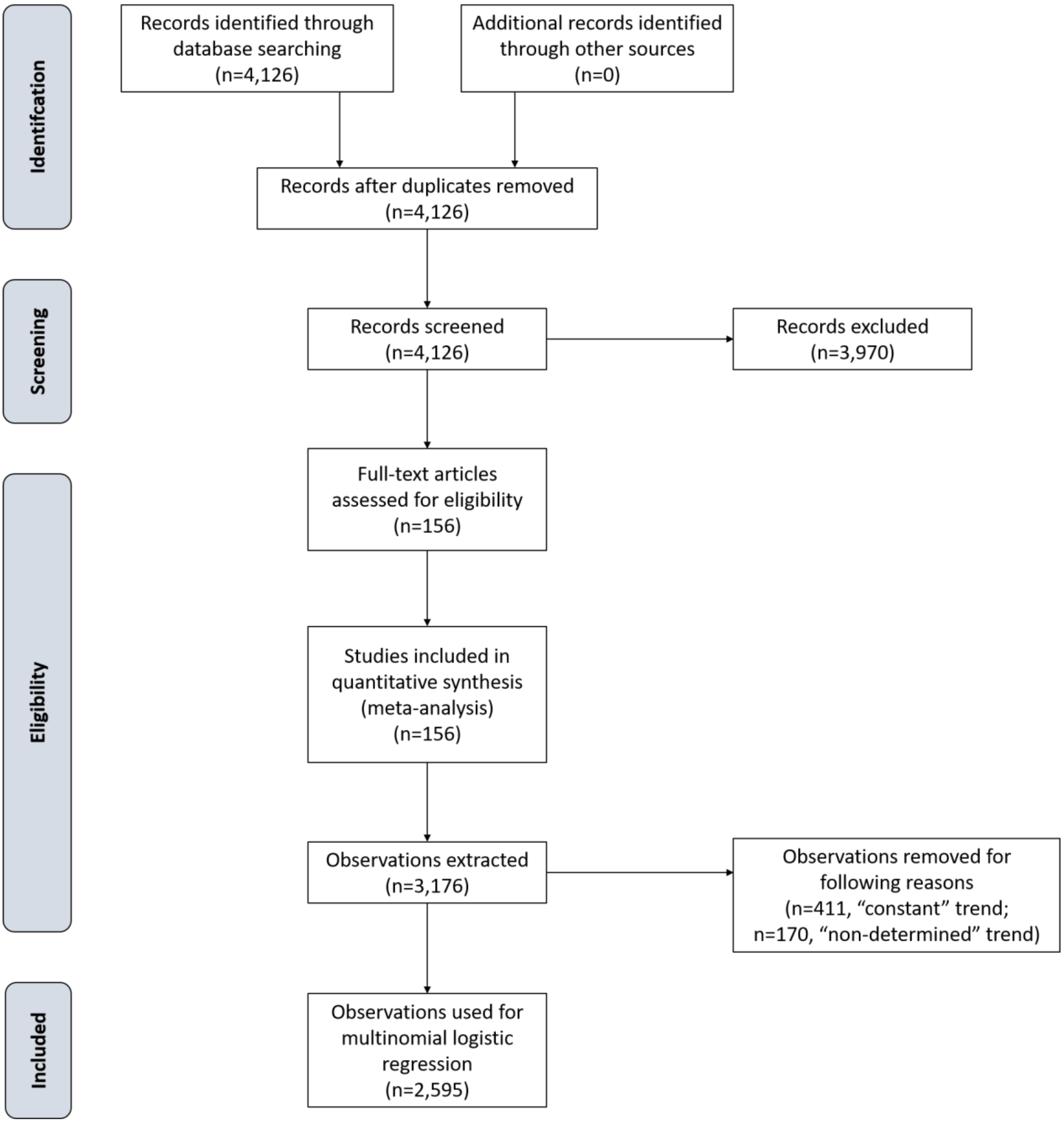
